# A Paradoxical Tumor Antigen Specific Response in the Liver

**DOI:** 10.1101/2024.09.19.614002

**Authors:** Rajiv Trehan, Xiao Bin Zhu, Patrick Huang, Xin Wang, Marlaine Soliman, Dillon Strepay, Amran Nur, Noemi Kedei, Martin Arhin, Shadin Ghabra, Francisco Rodríguez-Matos, Mohamed-Reda Benmebarek, Chi Ma, Firouzeh Korangy, Tim F. Greten

## Abstract

Functional tumor-specific CD8+ T cells are essential for an effective anti-tumor immune response and the efficacy of immune checkpoint inhibitor therapy. In comparison to other organ sites, we found higher numbers of tumor-specific CD8+ T cells in primary, metastatic liver tumors in murine tumor models. Despite their abundance, CD8+ T cells in the liver displayed an exhausted phenotype. Depletion of CD8+ T cells showed that liver tumor-reactive CD8+ T failed to control liver tumors but was effective against subcutaneous tumors. Similarly, analysis of single-cell RNA sequencing data from patients showed a higher frequency of exhausted tumor-reactive CD8+ T cells in liver metastasis compared to paired primary colon cancer. High-dimensional, multi-omic analysis combining proteomic CODEX and scRNA-seq data revealed enriched interaction of SPP1+ macrophages and CD8+ tumor-reactive T cells in profibrotic, alpha-SMA rich regions in the liver. Liver tumors grew less in Spp1^-/-^ mice and the tumor-specific CD8+ T cells were less exhausted. Differential pseudotime trajectory inference analysis revealed extrahepatic signaling promoting an intermediate cell (IC) population in the liver, characterized by co-expression of VISG4, CSF1R, CD163, TGF-βR, IL-6R, SPP1. scRNA-seq of a third data set of premetastatic adenocarcinoma showed that enrichment of this population may predict liver metastasis. Our data suggests a mechanism by which extrahepatic tumors facilitate the formation of liver metastasis by promoting an IC population inhibiting tumor-reactive CD8+ T cell function.

## Introduction

The liver is one of the most common sites for cancer metastasis. Up to 50% of cancer patients will present with or develop liver metastases during their course of disease.^1^ Unfortunately, the prognosis of patients with liver cancer metastasis is poor even after systemic therapy and resection.^1^ Moreover, patients with liver metastases receiving immune checkpoint inhibitors have significantly worse overall and progression-free survival than those without liver metastasis.^2^ This may be due to the liver’s unique immune environment favoring immune tolerance, which affects immunotherapy responses.^2^ Immune checkpoint inhibitor therapy aims to reinvigorate tumor-reactive CD8+ T cells, and new mechanistic insights drive novel treatment options.^3^

Tumor-reactive CD8+ T cells mediate anti-tumor immunity upon checkpoint inhibitor therapy and vaccines.^4,5^ Various mechanisms leading to CD8+ T cell dysfunction have been described. The mechanisms behind tumor-reactive CD8+ T cell dysfunction have been studied since defining in 1968 the Hellström paradox - the highly immunosuppressive tumor microenvironment (TME) must be overcome for effective immunotherapy.^4^ Similar to peripheral tolerance, tumor neoantigens early in tumor progression induce a hyporesponsive CD8+ T cell state.^4^ During late tumor progression, persistent antigen stimulation and the induced presence of immunosuppressive cells cause tumor-reactive CD8+ T cell dysfunction.^4^ Immune checkpoint inhibitor therapy aims to reinvigorate tumor-reactive CD8+ T cells, and new mechanistic insights drive novel treatment options.^5^

Numerous approaches for defining potentially tumor-reactive CD8+ T cells (pTRT) in single cell RNA-sequencing (scRNA-seq) data exist.^8–14^ Each approach tries to address the challenges of nonspecific classification of bystander T cells in tumors as well as the overrepresentation of exhausted pTRT clusters.^8–14^ Bystander T cell populations have been overrepresented despite the use of TCR clonality, application of TCR signaling signatures, and utilization of limited markers (CD39, PD1, Lag3, TCF7).^8–14^ Recent publications attempting to address these issues have instead used extensive gene signatures to profile tumor-reactive T cells across multiple tumor types, including colon cancer.^15,16^

T cell infiltration has been classically placed under the “hot and cold paradigm” where M2 macrophages play a major role in the immunosuppressive microenvironment of cold tumors.^17^ M2 tumor-associated macrophages (TAMs) have been associated with resistance to immunotherapy and have been a target for drug development.^17^ In contrast, bioinformatic analysis has postulated SPP1 macrophages to be distinct from the conventional M1 or M2 dogma, highlighting the need for in vivo data.^18^ However, there remains a lack of mechanistic insight into their specific role and function within the TME.^16^

Here, we apply a high-dimensional, integrated approach utilizing three human scRNA-seq data sets (with appropriate per-patient analysis), proteomic CODEX data, ex vivo systems, and murine models to identify an M2-independent, targetable, novel mechanism of tumor-reactive CD8+ T cell dysfunction specific to the liver. Additionally, we find that this mechanism provides a new paradigm for cancer cell metastasis to the liver through a unique IC population affecting profibrotic polarization.

## Results

### A paradoxical high frequency of tumor-antigen-specific (TAS) CD8+ T cells in tumor-bearing livers

The liver is a common site for metastasis from colon cancer. The frequency and phenotype of TAS CD8+ T cells were studied in mice bearing CT26 colorectal carcinoma using the well-established MHC class I tetramer loaded with AH1 peptide.^25,26^ CT26 cells were implanted into the liver to form intrahepatic (IH) tumors and into the skin to form subcutaneous (SQ) tumors (Figure 1A). First, we investigated the kinetics of TAS CD8+ T cells 7, 14 and 21 days post tumor injection in mice with intra-hepatic or subcutaneous CT26 tumors (Figure 1A). The frequencies of TAS CD8+ T cells in tumor, spleen, and tumor-free liver tissues were measured. Time course analysis revealed a marked early increase of TAS CD8+ T cells (∼10%) in intrahepatic tumor-infiltrating lymphocytes (TIL) one week after tumor injection compared to minimal levels of TAS CD8+ T cells in other tested tissues. The relatively higher frequency of TAS CD8+ T cells in intrahepatic TILs compared to subcutaneous TILs persisted throughout the entire observation period, peaking on day 14 (Figure 1B). The number of TAS CD8+ T cells increased over time in intrahepatic tumor-infiltrating lymphocytes (TIL) (Figure 1B). On day 14, the frequency of TAS CD8+ T cells was higher in the intrahepatic TIL compared to subcutaneous TIL (p = 0.0021), whereas no significant differences in tumor weights between intrahepatic and subcutaneous CT26 tumors were found at this time point, ruling out the contribution of tumor burden (p = 0.3792) (Figure 1B). Similar trends were seen when analyzing absolute counts (cells/gram tissue) on days 14 and 21 (Figures S1A-D). Interestingly, the tumor-free liver tissue surrounding the intrahepatic tumors also showed an increase in the frequency of TAS CD8+ T cells over time (Figure 1B). The frequency of TAS CD8+ T cells was lower in the spleen of mice with hepatic tumors as well as subcutaneous tumors. Both tumor-free mice and subcutaneous tumor-bearing mice showed consistently low frequency of TAS T cells in the liver across all analyzed time points (Figure 1B). Strikingly, the intrahepatic CT26 tumors grew at a faster rate than their subcutaneous counterparts despite the presence of a high number of tumor-infiltrating TAS CD8+ T cells (Figure 1C and D).^27^

**Figure 1:**
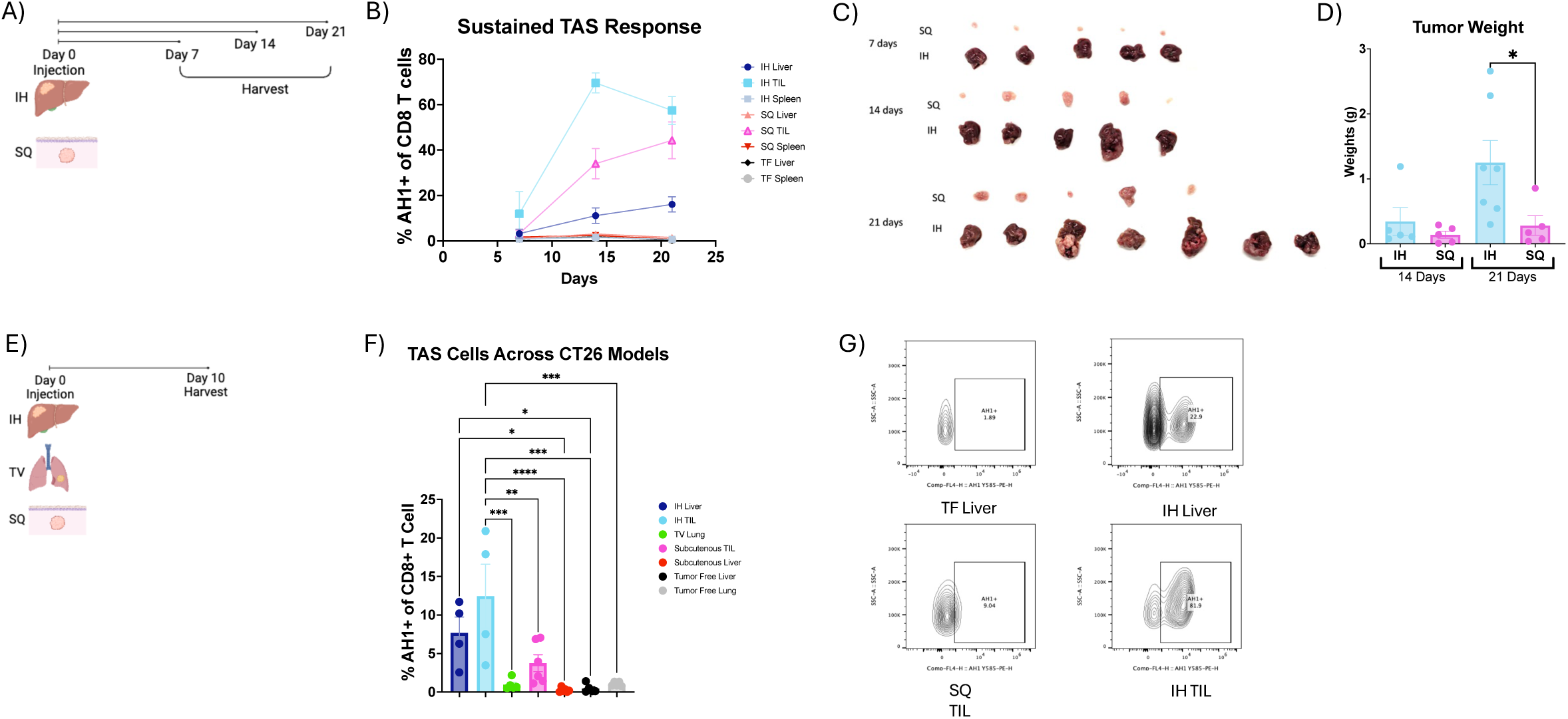
Tumor Antigen Specific (TAS) CD8 T Cells Across Murine Models. **A)** Experimental schematic for B-D. **B)** Frequency of AH1+ CD8+ T cells in liver and subcutaneous tumor-bearing mice compared to tumor free mice at days 7 (SQ, n=5; IH, n=5), 14 (SQ, n=5; IH, n=5) and 21(SQ, n=5; IH, n=7). **C)** Photo showing excised subcutaneous and intrahepatic tumors across days 7 (IH, n=5; SQ, n=5; TF, n=5), 14 (IH, n=5; SQ, n=5; TF, n=5) and 21 (IH, n=7; SQ, n=5; TF, n=5). **D)** Tumor weights at day 14 and 21 (IH, n=5; SQ, n=5;). Mice were euthanized at 3 weeks when tumors had reached humane clinical endpoints and, at day 21, the intrahepatic tumors had an average weight of 1.2495g while subcutaneous tumors weighed 0.28186g (p=0.0452). **E)** Experimental schematic for F and G. **F)** Frequency of AH1+ CD8+ T cells in various organs; IH, n=4; TV, n = 5; SQ, n=6; TF, n=5. **G)** Representative flow cytometry plots of F. Data was analyzed using ordinary one-way ANOVA for D and F.

Next, we included lung, another common organ to form metastasis. CT26 cells were implanted into the liver to form intrahepatic (IH) tumors, into the skin to form subcutaneous (SQ) tumors or injected into the tail vein (TV) to form intrapulmonary tumors (Figure 1E). Tumor, liver, lung and spleen tissue were harvested on day 10 for flow cytometric analysis (Figure 1E & S1E). The highest number of TAS CD8+ T cells (12.45%) was seen in tumor-infiltrating (TIL) CD8+ T cells in intrahepatic tumors (Figures 1F and G). The portal lymph node from liver tumor-bearing mice also showed a higher frequency of TAS CD8+ T cells compared to hilar and cutaneous lymph nodes (Figure S1F). A lower frequency (0.455%) of TAS CD8+ T cells was found in the spleen of the intrahepatic model (Figure S1F). In contrast to tumor-bearing lungs, the subcutaneous TIL had a high frequency (3.75%) of TAS CD8+ T cells (Figure 1F). Furthermore, lower numbers of TAS CD8+ T cells were found in the livers of mice with subcutaneous and pulmonary tumors (Figures S1G and H). To confirm the finding, TAS CD8+ T cells were also tested in the second model with B16F10 melanoma using an MHC class I tetramer loaded with TRP2 peptide, and similar results were found(Figures S1 I-L).^25,26^ In summary, tumors growing in the liver resulted in the highest frequency of TAS CD8+ T cells.

### Phenotype and function of TAS CD8+ T cells

To characterize the TAS CD8+ T cells in the intrahepatic and subcutaneous tumors, we characterized the phenotype of TAS CD8+ T cells by flow cytometry using markers of cell exhaustion (CD39, TIM-3, PD-1), cytotoxicity (Granzyme B), activation (CD69, PD-1) and proliferation (Ki67) (Figure S2).^28–30^ We also compared the AH1+ (TAS) CD8+ T cells to the AH1_neg_ CD8+ T cells which included the bystander CD8+ T cell population over time (Figure 1G).

We found the highest frequency of CD39+ TAS CD8+ T cells over time in the TILs and livers of mice with intrahepatic tumors. On day 14, intrahepatic TILs had a significantly higher frequency of CD39+ TAS T cells compared to subcutaneous TILs (p = 0.0287) which lasted till day 21 (p = 0.013). By day 21, the majority (89.7%) of the intrahepatic TILs expressed CD39 (Figure 2A).^28–30^ Notably, both livers from subcutaneous tumor-bearing mice and spleens from intrahepatic tumor-bearing mice showed an increase in the frequency of CD39+ TAS CD8+ T cells on day 21 (Figure 2A). We also found a correlation between the frequency of AH1+ (TAS) CD8+ T cells and CD39+ TAS CD8+ T cells in the liver and TIL of intrahepatic tumor-bearing mice (Figure 2B). This indicated that a higher frequency of TAS CD8+ T cells was correlated with an increase in the frequency of T cells with an exhausted phenotype in both the liver and tumor following intrahepatic injection. AH1+ (TAS) CD8+ T cells showed an approximate four-fold increase in CD39+ expression in the liver and a two-fold increase in the TIL of liver tumor-bearing mice compared to AH1_neg_ CD8+ T cells (Figure 2C). Similar results were found for TIM-3 expression (Figures 2D-F). Next, we studied Gzmb expression on CD8+ T cells.

**Figure 2.**
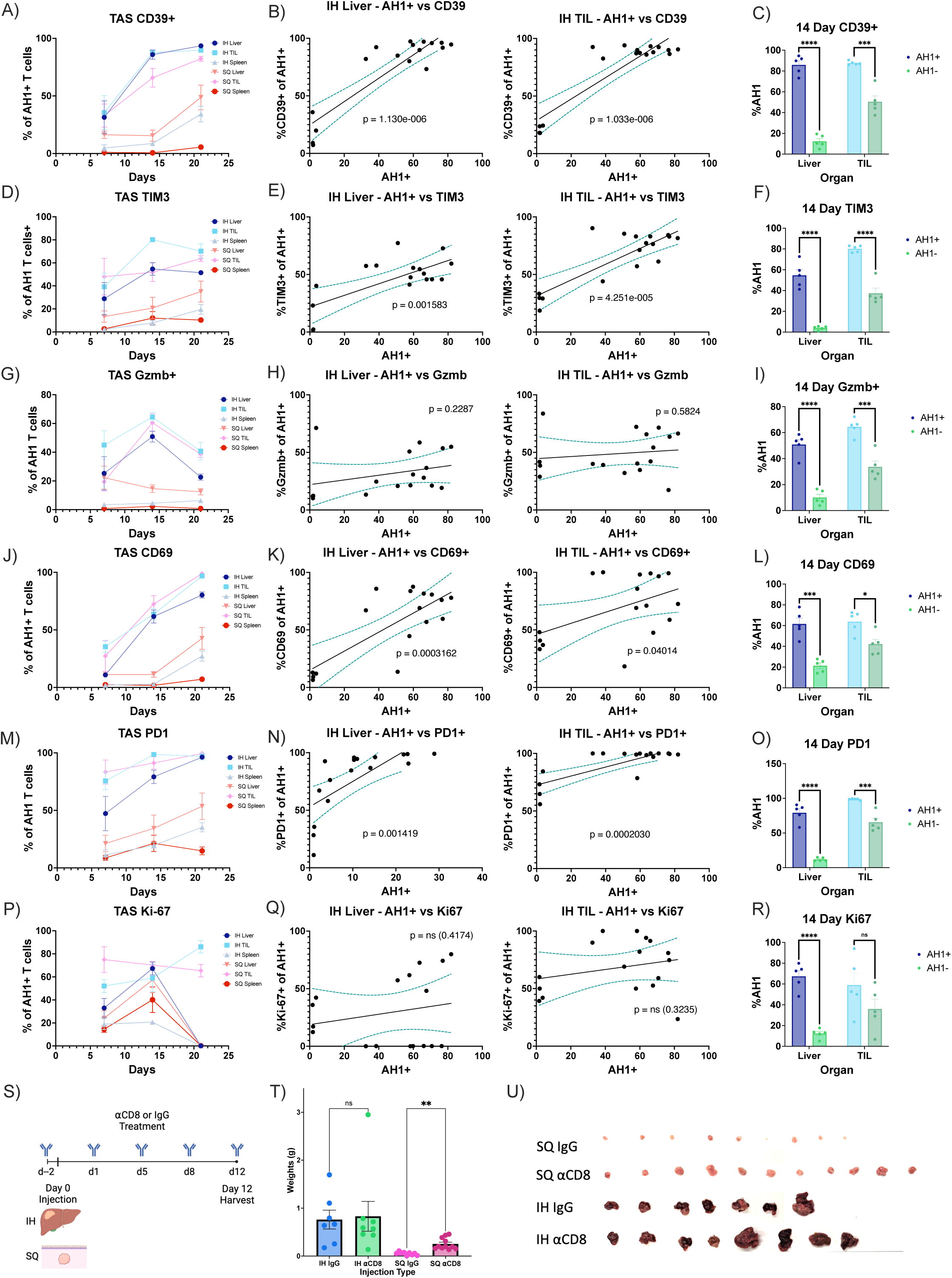
TAS CD8+ T cell phenotype and function. Frequency of **A)** CD39+, **D)** TIM-3+, **G)** Granzyme B+, **J)** CD69+, **M)** PD-1+, **P)** Ki-67+ among TAS CD8+ T cells in the TIL, liver and spleen of liver and subcutaneous tumor-bearing mice at days 7 (SQ, n=5; IH, n=5), 14 (SQ, n=5; IH, n=5) and 21(SQ, n=5; IH, n=7). Correlation between frequency of **B)** CD39+, **E)** TIM-3+, **H)** Granzyme B+, **K)** CD69+, **N)** PD-1+, **Q)** Ki-67+ AH1+ CD8+ T cells and frequency of AH1+ CD8+ T cells in the liver (left) and TIL (right) of liver tumor-bearing mice. Comparison of **C)** CD39+, **F)** TIM-3+, **I)** Granzyme B+, **L)** CD69+, **O)** PD-1+, **R)** Ki-67+ AH1+ to AH1-CD8+ T cells in the liver (left, n=17) and TIL (right, n=17) at day 14. **S)** Experimental schematic for T and U. **T)** Tumor weights after either IgG or αCD8 T cell antibody treatment after intrahepatic (IgG, n=7; αCD8, n=8) or subcutenous injection(IgG, n=9; αCD8, n=1) of tumor. **U)** Photo showing excised subcutaneous and intrahepatic tumors after either IgG or αCD8 treatment. Data was analyzed with simple linear regression (B,E,H,K,N,Q), unpaired t-tests (C,F,I,L,O,R) and brown-forsythe and welch anova (T,U).

The expression of the cytotoxic marker Gzmb in TAS CD8+ T cells showed an overall pattern of initial increase followed by a sharp drop following tumor progression. The frequency of Gzmb+ TAS CD8+ T cells both in the intrahepatic TILs and surrounding liver tissue, as well as subcutaneous TILs increased at day 14 but drastically decreased by day 21 (Figures 2G-I). In contrast to our findings regarding the exhaustion phenotype, no significant correlation was observed between the frequency of AH1+ (TAS) CD8+ T cells and the frequency of Gzmb+ TAS CD8+ T cells in the intrahepatic liver and TIL (Figure 2H). Thus, we found no correlation between a stronger TAS response and cytotoxicity. On day 14, AH1+ CD8+ T cells expressed increased levels of Gzmb compared to AH1_neg_ cells both in the intrahepatic TILs and the surrounding hepatic tissue of tumor-bearing livers (Figure 2I).

There was a consistently high frequency (96.84% on day 21) of CD69+ TAS CD8+ T cells over time in the TILs and livers of mice with liver tumors as well as in the TILs of mice with subcutaneous tumors (Figure 2J). Similar to CD39, both the livers from mice with subcutaneous tumors and the spleens from mice with hepatic tumors showed an unexpected increase in the frequency of CD69+ TAS CD8+ T cells on day 21 (Figure 2J). There was a correlation between the frequency of AH1+ (TAS) CD8+ T cells and the frequency of CD69+ TAS CD8+ T cells in both the liver and TIL after intrahepatic injection (Figure 2K). This points to an activated phenotype in TAS CD8+ T cells in both the liver and tumor after intrahepatic injection. The frequency of CD69+ AH1+ (TAS) CD8+ T cells was higher than AH1_neg_ CD8+ T cells in both the liver and the TIL of liver tumor-bearing mice (Figure 2L). Similar results were found for PD-1 (Figure 2M-O).^31^

There was no discernible pattern in the expression of Ki67+ in TAS CD8+ T cells over time within the intrahepatic and subcutaneous TILs (Figure 2P). Moreover, no correlation was seen between the frequency of AH1+ (TAS) CD8+ T cells and Ki67+ TAS CD8+ T cells in the liver and TIL following intrahepatic injection (Figure 2Q) indicating that the higher frequency of TAS CD8+ T cells was not due to an increase in proliferation. AH1+ (TAS) CD8+ T cells showed an approximately three-fold higher Ki67+ frequency in the liver with no difference in the TIL of liver tumor-bearing mice compared to AH1_neg_ CD8+ T cells (Figure 2R).

Next, we evaluated the effect of CD8+ T cell depletion on the growth of intrahepatic and subcutaneous tumors (Figure 2S). CD8+ T cell depletion using an anti-CD8α antibody was confirmed by flow cytometry (Figure S3B). Depletion of CD8 T cells did not affect the growth of intrahepatic tumors while subcutaneous tumors were significantly larger (p=0.0011) (Figures 2T, U; S3A). Thus, we conclude that dysfunctional CD8+ T cells in the liver failed to control intra-hepatic tumor growth but effectively controlled the growth of subcutaneous tumors.

In summary, these findings indicate that TAS CD8+ T cells showed an exhausted and dysfunctional phenotype in mice with liver tumors. Next, we decided to study human TAS CD8+ T cells within and outside the liver, as well as the interaction patterns of the intrahepatic TAS CD8+ T cells with other cells in the tumor microenvironment.

### SPP1+ macrophages interact with pTRT cells in humans

Given our findings from the mouse studies, we decided to investigate human tumor-reactive CD8+ T cells using publicly available processed and annotated scRNA-seq data from 10 patients with primary colorectal cancer tumor (PT), surrounding colonic tissue (PN), metastatic intrahepatic tumor (MT), surrounding hepatic tissue (MN), tumor-draining lymph nodes (LN), and peripheral blood mononuclear cells (PBMC) (Figure 3A).^32^ We obtained and validated the previously published cluster annotations for this data.^32^ A total of 304,816 cells among 78 clusters with published annotations were obtained from which 26 represented myeloid clusters (Figure S5A-B).^32^ Additionally, 14 CD8+ T cell clusters (previously published) were obtained including an (1) exhausted cluster expressing LAYN and LAG3, (2) tissue-resident memory cluster expressing KLRB1 and KLRD1, (3) CD160 IEL cluster expressing KLRC2, (4) HSPA1A cluster expressing HSPA6 and HSPA1B and a (5) proliferative cluster expressing MKI67 (Figure S5C).^32^

**Figure 3.**
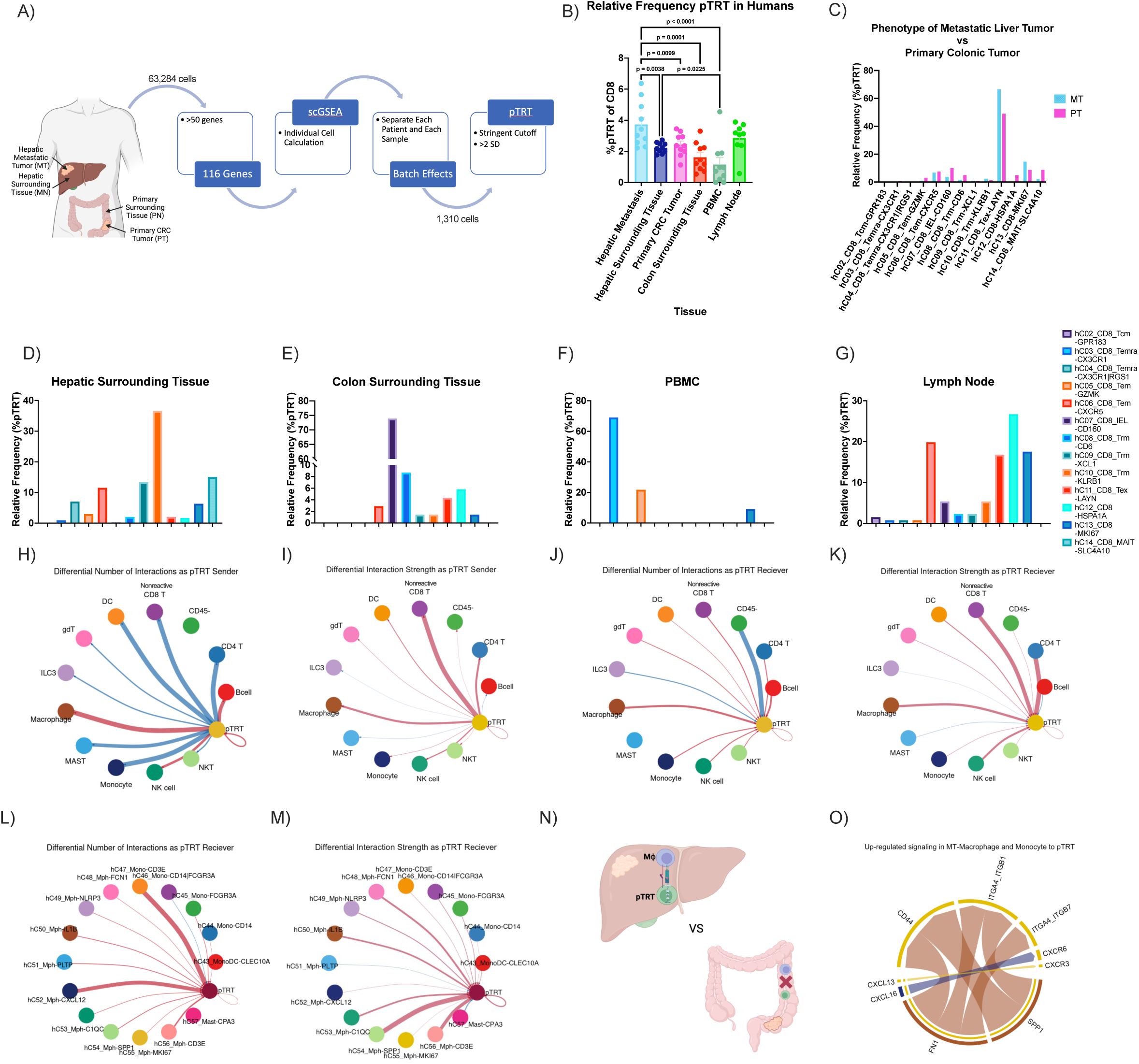
scRNA-seq analysis of human tumor-reactive CD8+ T cells. **A)** Schematic of scRNA-seq analysis to define pTRT CD8+ T cells using scGSEA. **B)** Frequency of pTRT CD8+ T cells in various organs; hepatic metastasis, n= 10; hepatic surrounding tissue, n=10; primary CRC tumor, n=10; colon surrounding tissue, n=10; PBMC, n=10; lymph node, n=9. Frequency of pTRT that are found in each CD8+ T cell cluster in **C)** hepatic metastisis (n=10) compared to primary CRC tumor (n=10), **D)** hepatic surrounding tissue (n=10), **E)** colon surrounding tissue (n=10), **F)** PBMC (n=10) and **G)** lymph node (n=9) with cluster labels found on the right of G. CellChat cellular communication analysis for communications that the pTRT are sending to all major cell types comparing hepatic metastasis to primary CRC tumors in the (**H**) number of interactions and (**I**) strength of interactions based on law of mass action. All interactions that were found to be statistically greater in the hepatic metastasis (n=10) are shown in red and in the primary CRC tumor (n=10) are shown in blue. Similarly, CellChat cellular communication analysis for communications that the pTRT are receiving from all major cell types comparing the (**J**) number of interactions and (**K**) strength of interactions between hepatic metastasis to primary CRC tumors. CellChat cellular communication analysis for communications that the pTRT are sending to macrophage clusters comparing hepatic metastasis to primary CRC tumors in the (**L**) number of interactions and (**M**) strength of interactions. **N)** Schematic of CellChat pathway analysis (**O**) comparing macrophage-pTRT CD8+ T cell interaction pathways upregulated in hepatic metastasis (n=10) compared to primary CRC tumor (n=10). Data were analyzed using paired one-way anova analysis (mixed effect analysis) (**B**) and a paired wilcox test (default for CellChat) (H-M,O).

We used single-cell Gene Set Enrichment Analysis (scGSEA) with more than 100 gene signatures to identify potentially tumor-reactive (pTRT) human CD8+ T cells (Figure 3A).^15,16^ We applied highly stringent cutoffs for the identification of pTRT cells to facilitate the identification of biologically valid mechanisms. CD8+ T cells (1,310 cells) that met this cutoff were labeled as pTRT cells and the rest of these cells were labeled as nonreactive CD8+ T cells hereafter (Figures S5D-F). As a negative control, scGSEA was performed using a randomly generated set of 500 genes which showed no differences across any organ sites (Figure S4B).

The highest frequency of pTRT CD8+ T cells among CD8+ T cells was found in TILs derived from liver metastasis followed by lymph nodes and primary CRC TILs (Figure 3B, S4A and 5F). Moreover, the frequency of intrahepatic pTRT CD8+ TILs was higher than that in the surrounding hepatic tissue (Figure 3B) analogous to findings in mice (Figure 1E). We also found a high pTRT frequency in tumor draining lymph nodes in the human data set (Figure 3B) analogous to our results in mice (Figure S1B).

We then determined the distribution of pTRT cells in previously identified T cell clusters for each tissue type (Figures 3C-G). Interestingly, pTRT cells were identified across a range of T cell clusters which allowed for study of both non-exhausted and exhausted pTRT cells. The majority (66.58%) of pTRT in intrahepatic TILs were found in the exhausted cluster (Tex-LAYN) (Figures S5D and E) similar to our murine models (Figures 2A and D). A greater proportion of the pTRT cells resided in the T exhausted cluster (LAYN) in intrahepatic TILs compared to primary CRC TILs (Figure 3C). The majority of pTRT in colonic tissue were CD160+ intraepithelial lymphocytes (IEL), while most of the pTRT in the surrounding hepatic tissue were tissue-resident memory (TRM) cells (Figures 3D and E). CD160+ IEL have been reported to show a cytotoxic function, particularly in the setting of viral diseases. Additionally, TRM T cells have been correlated to improved response to immunotherapy as these cells can be locally activated to provide anti-tumor function.^33–35^ The majority of pTRT cells in the lymph nodes were HSPA1A+ which is enriched in stressed T cells correlating to clinically nonresponsive tumors (Figure 3G).^36^ The lymph node pTRT cells also highly expressed proliferative marker MKI67 (Figure 3G).

We next analyzed cell-cell communication networks that are enriched in the intrahepatic tumors compared to primary CRC tumors using CellChat analysis. These networks can be measured either in number or by strength which is founded in the law of mass action.^37^ All CellChat analysis was normalized for the frequency of cell types when creating the initial CellChat object. We analyzed aggregated cell-cell communication networks specifically to and from pTRT cells that were enriched in the intrahepatic TILs, represented by red connection vectors, as compared to primary CRC TILs, represented by blue connection vectors (Figures 3H-K). Interactions between nonreactive CD8+ T cells to pTRT CD8+ T cells indicate an expected shared overlap between all CD8+ T cells. When analyzing broad cluster labeling, we were interested in identifying cell interactions highlighted in red (increased in liver metastasis compared to primary CRC) that were consistently sending and receiving signals from pTRT as characterized both by number and strength (Figures 3H-K). We observed that pTRT showed a consistent pattern of sending and receiving signals to and from macrophages more in intrahepatic TILs than in primary CRC TILs in both frequency and strength (Figures 3 H-K). Furthermore, the same cell-cell interactions between macrophages and pTRT cells were more prominent in the metastatic tumor compared to the surrounding hepatic tissue (Figures S6A and B). We also found that the macrophage and pTRT cell-cell interaction was more pronounced in the surrounding hepatic tissue as compared to the surrounding colonic tissue (Figures S6C and D). Next, we studied macrophages in greater detail. When analyzing cluster labeling for macrophages at a granular level, pTRT cells showed increased communication to CXCL12 macrophages and CD3E monocyte clusters in intrahepatic TILS compared to primary CRC TILs (Figure 3L). We also observed that pTRT received increased communication from SPP1 as well as CD3E macrophage clusters (Figure 3M). Next, we analyzed the cell-cell communication network from all macrophages to pTRT cells to identify pathways enriched in intrahepatic TILs as compared to primary CRC TILs (Figure 3N). Across all macrophage clusters, SPP1 and FN1 ligands from macrophages interacted with CD44, as well as integrin receptors ITGA4_ITGB1 and ITGA4_ITGB7, on pTRT cells to a greater degree in the intrahepatic TILs as compared to primary CRC TILs (Figure 3O).

In summary, a high frequency of pTRT CD8+ T cells was found in patients with hepatic metastasis. Additionally, intrahepatic pTRT cells preferentially interacted with SPP1+ and FN1+ macrophages. Previous studies have found SPP1+ macrophages to correlate with poor clinical outcomes in various cancer types.^18,38,39^ SPP1+ expression in healthy liver has been reported to be minimal and mainly localized to Kupffer cells compared to other immune cells.^40^ Fibronectin is a critical component of the extracellular matrix in liver fibrosis and has been linked to tumor recurrence after curative treatment.^41,42^ Given these enriched interactions, the role of SPP1+ and FN1+ macrophages in TAS CD8+ T cell dysfunction was further studied.

### SPP1+ macrophages cause pTRT cell dysfunction

Previous human scRNA-seq analyses have shown that SPP1+ macrophages represent a larger proportion of macrophages in liver metastasis compared to primary colonic tumors.^32^ We next profiled the macrophages in our murine models to see whether an enriched SPP1 and FN1 expressing macrophage population was also found in murine hepatic tumors. Mice were injected with CT26 tumor cells either intrahepatically or subcutaneously then intrahepatic and subcutaneous tumors were harvested after 15 days (Figure 4A) and TILS were analyzed by flow cytometry (Figure S7). There was a significantly higher frequency of Spp1+ macrophages in intrahepatic TILs compared to subcutaneous TILs (Figure 4B). Additionally, we found a significantly higher expression of PD-L1 on Spp1+ macrophages in the intrahepatic TILs compared to subcutaneous TILs (Figure 4C). These Spp1+ macrophages were distinct from M2 phenotype (CD163+ CD206+) which have protumor function and showed significantly decreased expression in the intrahepatic TILs as compared to subcutaneous TILs (Figure 4D). Finally, the expression of Spp1 by macrophages showed a significant correlation with tumor weight (Figure 4E). In summary, similar to the human scRNA-seq results, an enriched Spp1 macrophage population was seen in intrahepatic TILs which correlated with tumor size and showed a distinct phenotype from the M2 macrophages.

**Figure 4.**
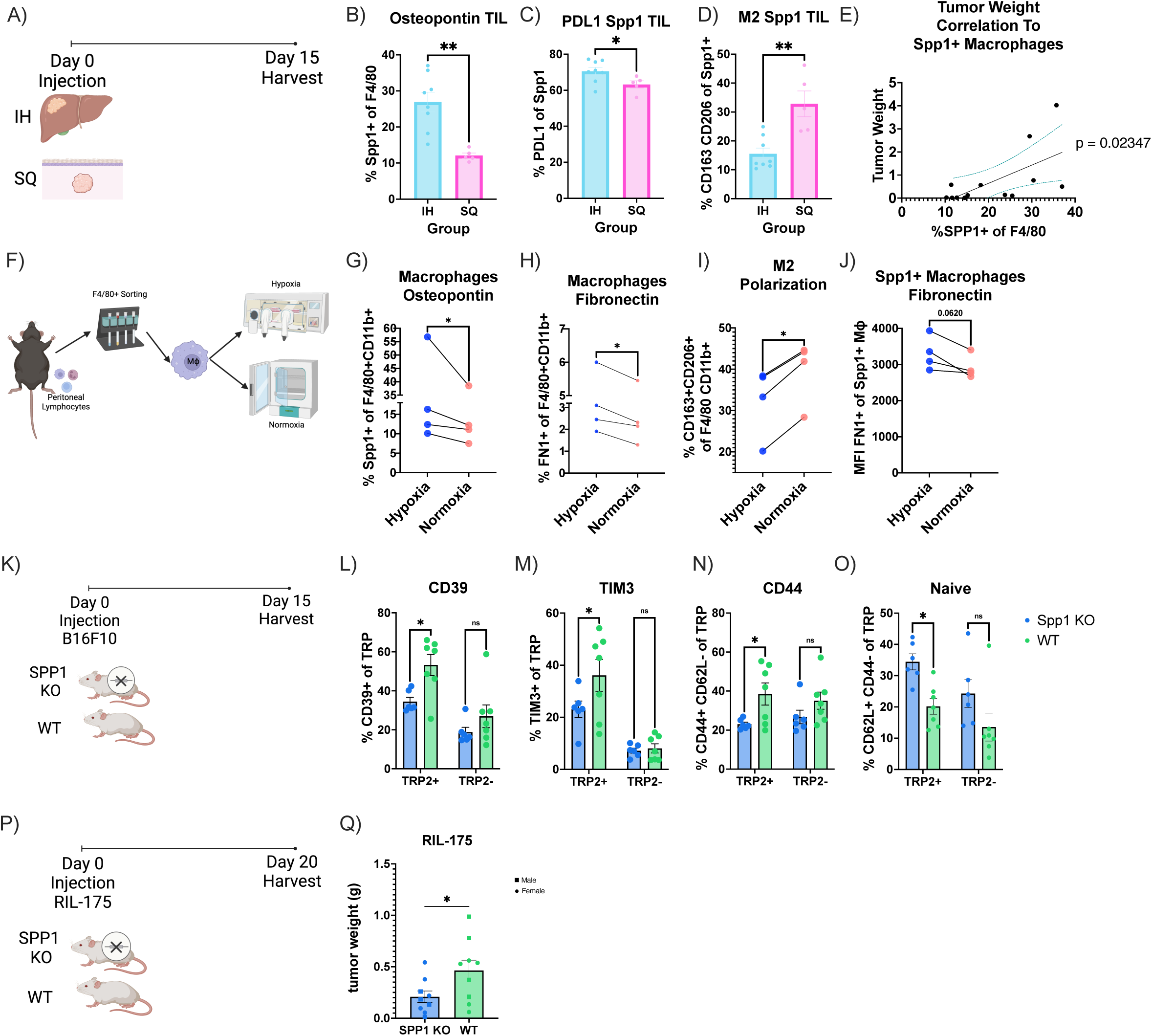
Spp1+ Macrophages cause TAS CD8+ T cell dysfunction. **A)** Experimental schematic for B-E. **B)** Frequency of Spp1+ macrophages in intrahepatic (n=8) and subcutenous TILs (n=5). **C)** PDL1+ frequency of Spp1+ macrophages in intrahepatic (n=8) and subcutenous TILs (n=5). **D)** CD163+ CD206+ (M2) frequency of Spp1+ macrophages in intrahepatic (n=8) and subcutenous TILs (n=5). **E)** Correlation between orthtopic tumors and frequency of Spp1+ macrophages (n=13). **F)** Experimental schematic for G-J. **G)** Frequency of SPP1+ macrophages after hypoxia and normoxia (n=4). **H)** FN1+ frequency of macrophages after hypoxia and normoxia (n=4). **I)** CD163+ CD206+ (M2) frequency of macrophages after hypoxia and normoxia (n=4). **J)** Spp1+ macrophage expression of FN1 (MFI) after hypoxia and normoxia (n=4). **K)** Experimental schematic for L-O. **L)** Frequency of CD39+ TRP2+ (TAS) CD8+ T cells (left) and CD39+ TRP2-(TAS) CD8+ T cells (right) in Spp1 KO compared to WT mice. **M)** Frequency of TIM-3+ TRP2+ CD8+ T cells (left) and TIM-3+ TRP2-CD8+ T cells (right) in Spp1 KO (n=6) compared to WT mice (n=7). **N)** Frequency of CD44+ CD62L-TRP2+ CD8+ T cells (left) and CD44+ CD62L-TRP2-CD8+ T cells (right) in Spp1 KO (n=6) compared to WT mice (n=7). **O)** Frequency of CD44-CD62L+ TRP2+ CD8+ T cells (left) and CD44-CD62L+ TRP2-CD8+ T cells (right) in Spp1 KO (n=6) compared to WT mice (n=7). **P)** Experimental schematic for Q. **Q)** RIL-175 tumor weight comparing in Spp1 KO (n=5) compared to WT mice (n=5). Data was analyzed using ordinary one way Anova (B-D), paired t-test (G-J,M,T,V), simple linear regression (E), 2 way anova (L-O).

Previous studies have shown a correlation between hypoxia and SPP1+ macrophages.^18^ Therefore, we utilized hypoxic conditions to induce a Spp1 positive phenotype in macrophages. Peritoneal macrophages were isolated (Figure 4F, S8 and B)^43^ and placed under normoxic or hypoxic (0.5% O_2_) conditions for 24 hours (Figure 4F). Afterward, the cells were analyzed by flow cytometry. Under hypoxic conditions, macrophages expressed significantly higher levels of Spp1 (Spp1^high^) compared to normoxic (Spp1^low^) conditions (Figure 4G, S8C). Additionally, under hypoxic conditions, macrophages had a significant increase in FN1 expression compared to normoxic conditions (Figure 4H). Interestingly, similar to our findings in murine models, the expression of M2 markers (CD163+ CD206+) on macrophages decreased under hypoxic conditions (Figure 4I). SPP1+ macrophages expressed higher levels of FN1 under hypoxia indicating co-expression of Spp1 and FN1 (Figure 4J).

To determine the effect of Spp1+ macrophages *in vivo*, we implanted intrahepatic B16F10 tumors in both homozygous Spp1 knockout (KO) and wildtype (WT) mice (Figure 4K). Flow cytometric analysis was performed on day 15. We found that TAS CD8+ T cells from Spp1 KO mice had decreased expression of exhaustion markers, CD39 and TIM-3 as compared to WT mice (Figures 4L and M). Moreover, this difference was only observed in TAS (TRP2+) CD8+ T cells and not in TRP2-CD8+ T cells (Figures 4L and M). CellChat analyses of human T cells and macrophages showed an interaction of SPP1 on macrophages with CD44 on pTRT CD8+ T cells (Figure 3O) leading us to investigate whether a similar observation could be made in mice. Compared to WT mice, CD44+ CD62L-expression was also decreased on TAS CD8+ T cells while the naïve (CD44-CD62L+) phenotype was increased in Spp1 KO mice (Figures 4N and O). Similar to the exhaustion markers, these changes were significant only for TRP2+ CD8+ T cells rather than TRP2-C8+ T cells (Figures 4Q and R). Finally, to examine the effect of SPP1 on overall tumor burden, we injected a mouse HCC cell lines (RIL-175) orthotopically into the livers of Spp1 KO and WT mice (Figure 4P). Mice were sacrificed after 20 days (RIL-175) post-injection and tumor weights were measured. Knocking down Spp1 significantly decreased tumor sizes with RIL-175 (Figure 4Q).

In summary, our findings show that SPP1-CD44 interactions were enriched in the liver of stage IVA+ CRC patients based on scRNA-seq analysis. Subsequently, we found that Spp1+ macrophages promote TAS CD8+ T cell exhaustion and dysfunction affecting tumor burden. Additionally, we showed Spp1+ macrophages to co-express FN1 (Figures 4H and J). To better understand the effect of fibronectin co-expression, we next explored the role of extracellular matrix (ECM) changes in the liver TME.

### Macrophage and CD8+ T cell interactions are enriched in αSMA+ environments

Previous studies have shown SPP1+ macrophages interact with fibroblasts.^19,20^ However, based on the aforementioned cell chat analysis, we postulate macrophage induced differences in ECM components, including fibronectin, affect macrophage-CD8+ T cell interactions. We recently performed a spatial co-detection by indexing (CODEX) imaging analysis as well as scRNA-seq analysis from liver tumors and adjacent tissue of HCC patients.^44^ We repeated scGSEA on this HCC scRNA-seq data set with the same approach as our previous scRNA-seq analysis to define pTRT CD8+ T cells in these HCC patients (Figure 3A). CellChat analysis of our RNA-seq data confirmed that SPP1+ macrophage interactions with pTRT cells also occurred in HCC (Figure S9A).^44^ The CODEX imaging identified cell types at a single-cell resolution using our previously published thresholds to define the positive population for each marker (Figure 5A).^44^ This high-dimensional, integrated multi-omic analysis of protein (CODEX) and mRNA expression (scRNA-seq) connects intercellular signaling on a spatial and transcript level in the context of a single pathology.

**Figure 5.**
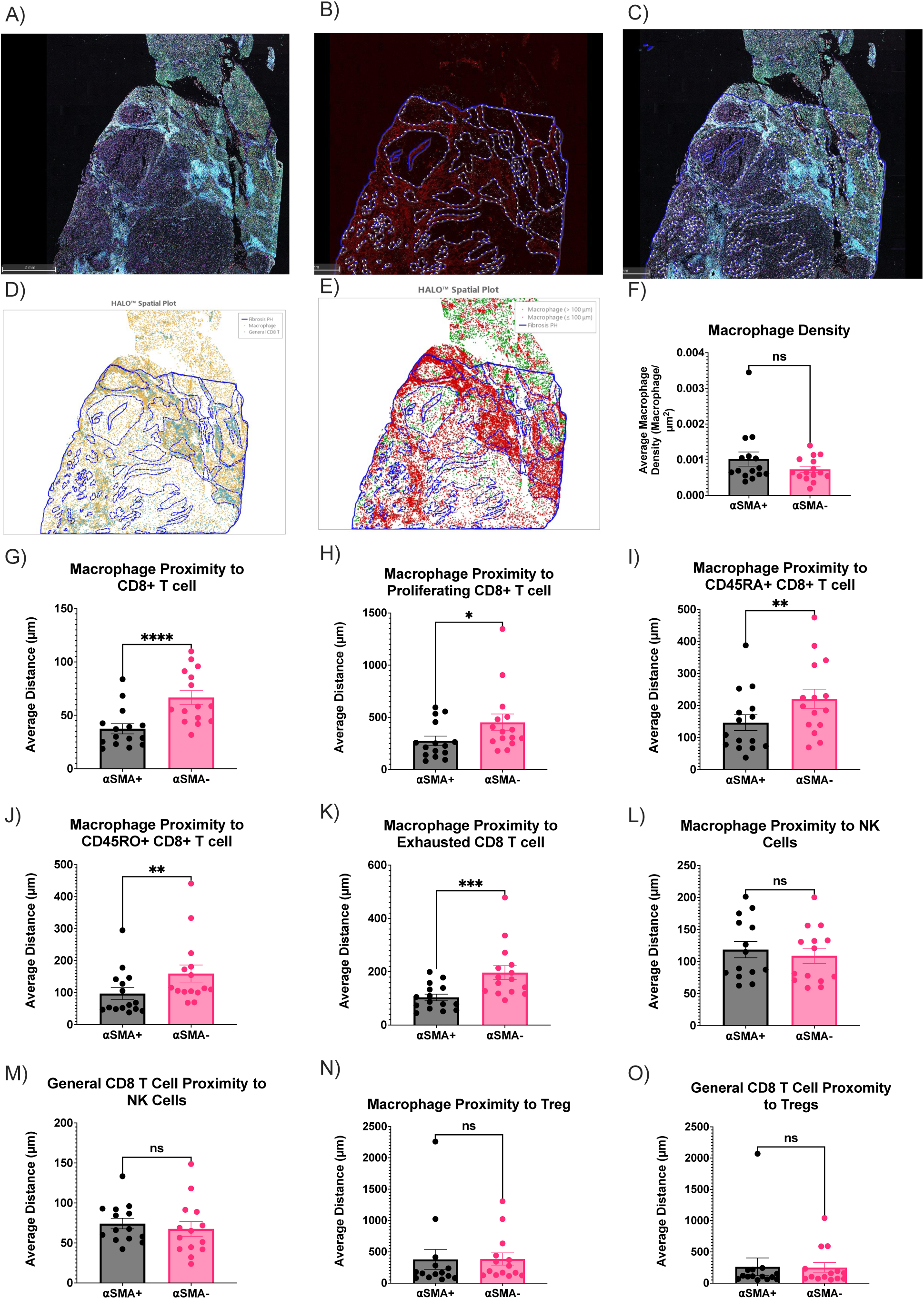
CODEX analysis of macrophage and CD8+ T cell interactions in αSMA rich regions. **A)** Overview of whole-tissue section of human HCC sample with 37-plex CODEX panel. **B)** Annotations (blue) of profibrotic αSMA+ (red) regions. **C)** Overlay of annotations (B) on whole-tissue section (A). **D)** Single cell resolution annotation of macrophages and CD8+ T cells. **E)** Representative spatial plot of macrophages proximity analysis to CD8+ T cells where macrophages within 100 µM of CD8+ T cells are labeled in red and macrophages farther than 100 µM of CD8+ T cells are labled in green. **F)** Macrophage density in αSMA+ and αSMA-regions (n=15). **G)** Macrophage proximity to CD8+ T cells in αSMA+ and αSMA-regions (n=15). **H)** Macrophage proximity to Ki67+ CD8+ T cells in αSMA+ and αSMA-regions (n=15). **I)** Macrophage proxmity to CD45RA+ CD8+ T cells in αSMA+ and αSMA-regions (n=15). **J)** Macrophage proximity to CD45RO+ CD8+ T cells in αSMA+ and αSMA-regions (n=15). **K)** Macrophage proxmity to CD39+ CD8+ T cells in αSMA+ and αSMA-regions (n=15). Data was analyzed by paired t-test (F-O). **L)** Macrophage proximity to NK cells in αSMA+ and αSMA-regions (n=15). **M)** CD8+ T cell proximity to NK cells in αSMA+ and αSMA-regions (n=15). **N)** Macrophage proximity to Treg in αSMA+ and αSMA-regions (n=14). **O)** CD8+ T cell proximity to Tregs in αSMA+ and αSMA-regions (n=14).

FN1 expression in the extracellular matrix mediates TGFβ induced fibroblast contraction and αSMA expression.^45^ Thus, we annotated our CODEX images for regions that were both rich in αSMA and morphologically overlapped with a profibrotic structure (Figures 5B and C). We then analyzed at a single cell resolution, CD8+ T cells and macrophages in profibrotic (αSMA+) regions compared to the surrounding tissue (αSMA-neg) regions (Figures 5D and E). Cell quantification analysis showed no difference between macrophage density between αSMA+ and αSMA-neg regions (Figure 5F). However, proximity analysis revealed that macrophages were closer to CD8+ T cells in αSMA+ compared to αSMA-neg regions (Figure 5K). Additionally, macrophages were closer to proliferating (Ki67+), CD45RA+, CD45RO+ and exhausted (CD39+) CD8+ T cells in the αSMA+ regions compared to αSMA-neg regions (Figures 5G-K). As a negative control for the next proximity analysis, there was no difference between the proximity of macrophages to NK cells between αSMA+ and αSMA-regions (Figure 5L). Additionally, no differences were observed between CD8+ T cells and NK cells within αSMA+ and αSMA-neg regions (Figure 5M). Similar results were found for Tregs (Figures 5N and O).

In summary, we investigated the role of profibrotic ECM changes in the liver TME by analyzing CODEX data of HCC patients. We found that CD8+ T cells including proliferating, CD45RA+, CD45RO+ and exhausted phenotypes were more likely to be in close proximity to macrophages in profibrotic αSMA+ regions than in αSMA-neg regions. We next aimed to reconcile the profibrotic interactions with the earlier flow cytometry results from the livers of mice with subcutaneous tumors.

### Profibrotic environment enriches SPP1

Thus far, both CODEX and CellChat results showed that macrophage-CD8+ T cell interactions were enriched in profibrotic, αSMA+ regions. In addition, time course experiments (Figure 6A) showed on day 21 an increase in CD39 (Figure 2A and 6B), TIM-3 (Figure 2D and 6C), PD1 (Figure 2M and 6D), CD69 (Figure 2J and 6E) expressing TAS CD8+ T cells in the liver of mice with subcutaneous tumors. We next asked whether increased profibrotic machinery in the liver could explain why there was such a high frequency of exhausted, activated TAS CD8+ T cells in the liver of mice with subcutaneous tumors.

**Figure 6.**
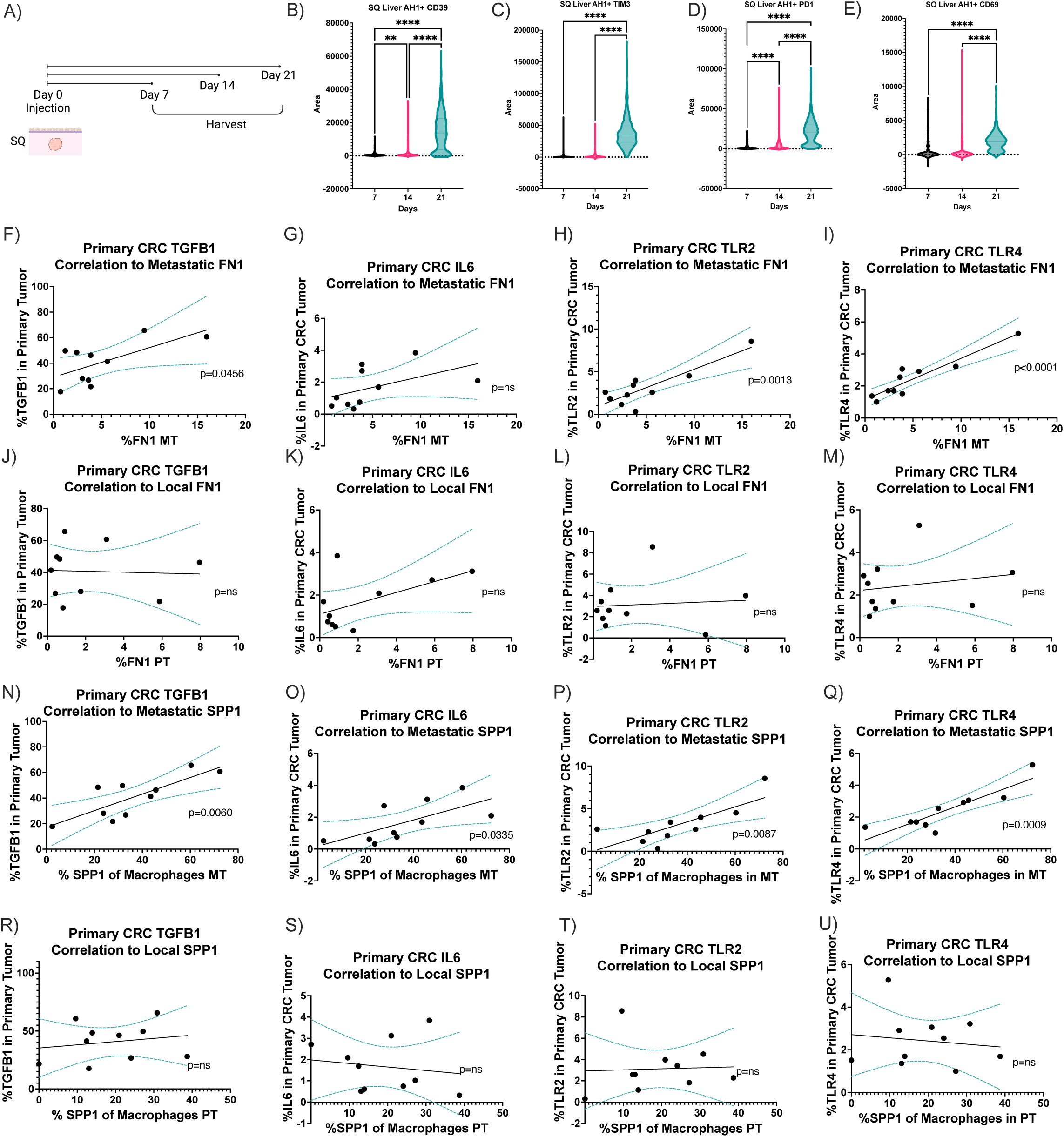
Exhausted, profibrotic polarization of the liver correlated to extrahepatic tumor signalling. **A)** Experimental schematic of B-E. AH1+ CD8+ T cells in the liver of subcutenous tumor bearing mice expressing CD39 **(A)**, TIM3 (**C**), PD1 (**D**), CD69 (**E**). Correlation of fibronectin expression in metastatic intrahepatic tumors (MT) to TGF-β **(F)**, IL-6 **(G)**, TLR-2 **(H)**, TLR-4 **(I)** in primary tumors (n=10). Correlation of fibronectin expression in primary CRC tumors (PT) to TGF-β **(J)**, IL-6 **(K)**, TLR-2 **(L)**, TLR-4 **(M) in** primary tumors (n=10). Correlation of SPP1 expression by macrophages in metastatic intrahepatic tumors (MT) to TGF-β **(N)**, IL-6 **(O)**, TLR-2 **(P)**, TLR-4 **(Q)** expression in primary tumors (n=10). Correlation of SPP1 expression by macrophages in primary CRC tumors to TGF-β **(R)**, IL-6 **(S)**, TLR-2 **(T)**, TLR-4 **(U)** in primary tumors (n=10). Data was analyzed was simple linear regression (F-U) and one-way anova (B-E).

TGF-β and IL-6 serve as important regulators of inflammation and fibrosis, and their production and action are potentiated by upstream TLR-2 and TLR-4.^46–51^ These are expressed on a wide variety of cells including hepatocytes, hepatic stellate cells, biliary epithelial cells, liver sinusoidal endothelial cells, dendritic cells and Kupffer cells.^46,47^

We questioned if the extrahepatic tumor could be upregulating profibrotic machinery in the liver. Additionally, we were interested in studying if the primary CRC tumor controls the liver TME in the same manner that it affects the local colonic TME. To answer this, we used human patient scRNA-seq data to correlate the intrahepatic SPP1 and FN1 response to profibrotic proteins in primary CRC tumors.

Interestingly, a significant correlation was found between the expression of TGF-β, IL-6, TLR-2 and TLR-4 by the primary CRC tumor and FN1 expression in the metastatic intrahepatic tumor (Figure 6F-I) but no correlation to the local primary CRC tumor (Figure 6J-M). Thus, extrahepatic tumor production of profibrotic proteins correlated with intrahepatic expression of fibronectin but no correlation to local fibronectin.

We next asked whether there was a correlation between SPP1 expression by macrophages in the profibrotic environments and TGF-β, IL-6, TLR-2 and TLR-4 expression levels in the primary CRC tumors of the patients. Interestingly, there was a significant correlation between TGF-β, IL-6, TLR-2 and TLR-4 in the primary CRC tumor and SPP1 expression by macrophages in the metastatic intrahepatic tumor (Figure 6N-Q). However, no such correlation to the local primary CRC tumor was found (Figure 6R-U). Additionally, no significant correlation was observed between M2 macrophage markers CD163 and CD206 in the intrahepatic tumor to TGF-β, IL-6, TLR-2 and TLR-4 in the primary CRC tumor (Figures S9B-I). In summary, extrahepatic tumor production of profibrotic proteins correlated with intrahepatic SPP1 expression but no correlation to intrahepatic M2 markers or local SPP1 expression.

In summary, we demonstrate a connection between TGF-β, IL-6, TLR-2 and TLR-4 expression by the primary CRC tumor to SPP1 and FN1 expression in the metastatic intrahepatic tumor. There was no correlation between the expression of these cytokines and the surrounding colonic microenvironment. We next analyzed the macrophage population to further elucidate the mechanism by which these cytokines preferentially affect the liver.

### Pseudotemporal Analysis Reveals a Liver-Specific Intermediate Cell Population

Using the previous annotated scRNA-seq data of paired CRC liver metastasis, 4,833 macrophage and monocyte cells in the liver were clustered (Figure 7A).^32^ Kupffer cell (KC) markers, VSIG4 and CSF1R, were found to be highly co-expressed in the top region of the UMAP (Figure S10A). We utilized pseudotime trajectory analysis of the intrahepatic macrophage and monocytes cells to infer the relationship of SPP1, TGF-βR and IL-6R to dynamic, continuous gene changes (Figure 7B).

**Figure 7.**
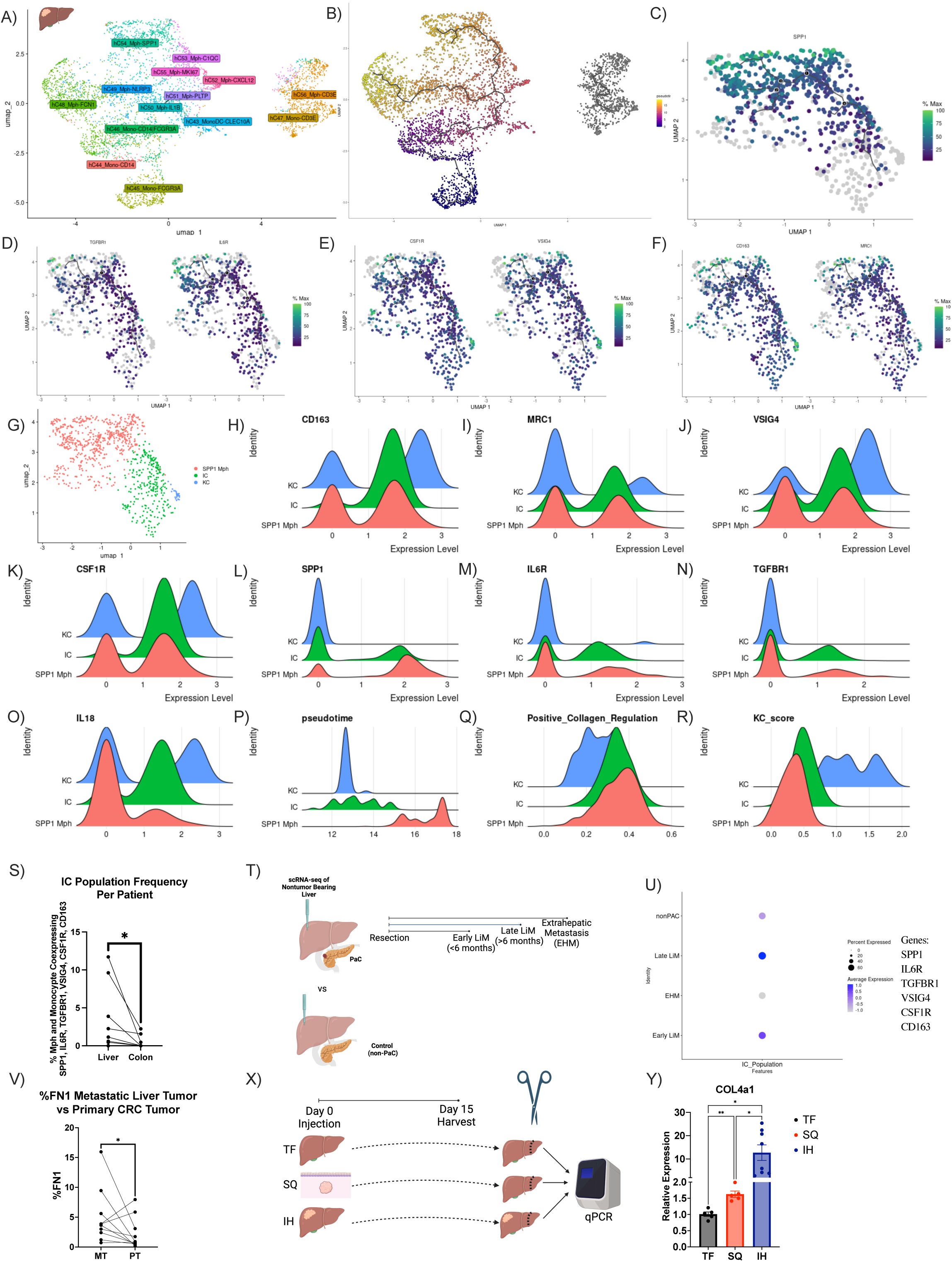
Intermediate cell population secondary to metastatic signalling. Intrahepatic macrophage and monocyte population UMAP from metastatic CRC tumor-bearing liver **(A)** (n=10). Pseudotime trajectory analysis (n=10) of intrahepatic macrophages and monocytes **(B)**. Top branch of pseudotime analysis showing correlation of SPP1 **(C)**, TGFBR1 and IL6R **(D),** CSF1R and VSIG4 (**E)** and CD163 and CD206 **(F)** with pseudotime trajectories (n=10). Subclustering of pseudotime branch related to SPP1 (n=10) **(G)** with ridge plot displaying expression of CD163 **(H)**, CD206 **(I)**, VSIG4 **(J)**, CSF1R **(K)**, SPP1 **(L)**, IL6R **(M)**, TGFBR1 **(N)**, IL18 **(O)**, pseudotime **(P)**, positive collagen regulation score **(Q)**, Kupfer cell (KC) score **(R)**. Frequency of cells coexpressing all markers of IC cluster in all macrophages and monocytes in the liver and the colon per patient (n=10) **(S)**. Experimental schematic **(T)** of scRNAseq analysis in U. Intermediate cluster score calculated in CD68+ per group **(U)**. **V)** Expression of fibronectin in metastatic intrahepatic tumor compared to primary CRC tumor. **X)** Experimental schematic of Y. **Y)** Collagen type IV subunit A qPCR expression in the liver of tumor free, subcutenous tumor and intrahepatic tumor-bearing mice. Data were analyzed was paired t-test (S,V) and one-way anova (Y).

Of all 24,622 genes tested, SPP1 was the 4^th^ highest correlation to a function of pseudotime (Moran’s I = 0.72) (Figure S10C). SPP1 trajectory analysis revealed differentiation from the branch point along the upper leaf of pseudotime (Figure 7C). Highest SPP1 expression was found at the terminal end of pseudotime trajectory analysis (Figure 7C). This leaf of pseudotime was analyzed for continual changes in the expression of relevant markers earlier in the trajectory (Figures 7D-F). TGF-βR1 and IL-6 were found to closely follow the trajectory and were expressed earlier in pseudotime as compared to SPP1 (Figure 7D). Similar trajectory results were found for KC markers CSF1R and VSIG4 (Figure 7E) as well as CD163 and CD206 (Figure 7F). Combining pathway analysis, scGSEA and trajectory analysis, genes that upregulate collagen production in the surrounding microenvironment were analyzed (Figure S10D). Collagen upregulation genes were expressed at an earlier pseudotime in the trajectory analysis as compared to SPP1. Throughout this analysis, we detected a portion of the cells on the UMAPs that co-expressed all these markers at an intermediate pseudotime. Thus, we next clustered all cells along this leaf of pseudotime and found 3 distinct populations (Figure 7G).

A KC population (Figure 7G) was found that highly expressed CD163 (Figure 7H), CD206 (Figure 7I), VSIG4 (Figure 7J), CSF1R (Figure 7K) and IL-18 (Figure 7O). This KC population also highly expressed previously published gene scores for identifying KCs (Figure 7R).^52^ The KC population had lower expression of SPP1 (Figure 7L), IL6R (Figure 7M), TGFBR1 (Figure 7N), positive collagen regulation (Figure 7Q). These cells were found at an earlier pseudotime (Figure 7P).

A cluster of macrophages (Figure 7G) highly expressed SPP1 (Figure 7L). This SPP1+ macrophage population downregulated CD163 (Figure 7H), CD206 (Figure 7I), VSIG4 (Figure 7J), CSF1R (Figure 7K), IL-18 (Figure 7O) and KC score (Figure 7R). This SPP1 macrophage population which distinctly clustered away from the KC population was found at the end of pseudotime (Figure 7P).

Recent literature identifies SPP1 macrophages as distinct from KCs in fatty liver.^24^ In contrast, in the liver TME, an intermediate cluster (IC) population (Figure 7G) was found to co-express intermediate levels of CD163 (Figure 7H), CD206 (Figure 7I), VSIG4 (Figure 7J), CSF1R (Figure 7K), SPP1 (Figure 7L), IL6R (Figure 7M), TGFBR1 (Figure 7N), IL18 (Figure 7O), and KC score (Figure 7R). This cluster had the highest expression of IL6R (Figure 7M) and TGFBR1 (Figure 7N) as well as positive collagen regulation (Figure 7Q). This cluster was isolated in an intermediate pseudotime state bridging the SPP1 and KC clusters (Figure 7P).

Performing a similar analysis for colonic macrophages and monocytes in paired CRC metastasis, 1,330 macrophage and monocyte cells were clustered (Figure S10E).^32^ Resident macrophage markers VSIG4 and CSF1R were found to be highly co-expressed in the left region of the UMAP (Figure S10B). We utilized pseudotime trajectory analysis of the intrahepatic macrophage and monocytes cells to infer the relationship of SPP1, TGF-βR and IL-6R to dynamic, continuous gene changes (Figure S10F). SPP1 was not correlated with pseudotime in the colon (Moran’s I = 0.29) (Figure S10G). Unlike the liver, no significant relationship was found between TGFBR1, IL6R, CSF1R, VSIG4, CD163, CD206 and pseudotime (Figures S10H-K). When comparing the frequency of cells co-expressing SPP1, TGFBR1, IL6R, CSF1R, VSIG4, CD163 per patient, the liver had a significantly larger IC population (Figure 7S). Bulk TCGA analysis with ssGSEA found that patients with enriched IC expression had worse survival in HCC but not in colon cancer (Figure S10L). In summary, an expanded IC population unique to the liver expressing high levels of TGFBR1 and IL-6R was found later in pseudotime compared to KCs.

To validate the above findings of a premetastatic niche in our murine models and liver metastasis, we next analyzed a third, recently published scRNA-seq data of patients with pancreatic adenocarcinoma (Figure 7T).^53^ Biopsies of the non-tumor bearing liver at the time of resection were analyzed using scRNA-seq (Figure 10M), and these patients were then followed to determine the incidence of liver metastasis. After QC, 26,627 total cells were plotted on a UMAP (Figure S10-M). 2 clusters containing 2,345 cells were found to express CD68 (Figure S10M). These clusters were then analyzed for IC scGSEA enrichment (Figure 7U). IC enrichment was detected in patients who would develop early and late liver metastasis (Figure 7U) as compared to control (non-pancreatic adenocarcinoma) or extrahepatic metastasis patients. Individually VSIG4, CSF1R and SPP1 were not highly predictive in patients who will develop liver metastasis in the future (Figure S10N). In summary, the IC population was enriched in the premetastatic niche which may predict future liver metastasis.

Given that the response to profibrotic factors is mainly observed in the liver rather than the primary CRC tumor, we next probed the fibronectin levels in the metastatic intrahepatic tumor in the human scRNA-seq data. The metastatic intrahepatic tumor had higher FN1 expression (Figure 6V). Thus far, we have shown a correlation between profibrotic proteins in the primary CRC tumor and the expression of FN1 and SPP1 in the metastatic intrahepatic tumor.

We next question whether mice with subcutaneous tumors also had changes in liver ECM components. Fibronectin is necessary for collagen matrix assembly and, while many types of collagen make up liver fibrosis ECM, collagen type 4 has been a valuable biomarker.^42,54^ Collagen type 4 levels were studied utilizing qPCR to investigate the effect of these profibrotic changes on the polarization of ECM components. This was chosen over H & E staining as no morphological changes in the liver architecture were expected. Subcutaneous tumor-bearing mice as well as liver tumor-bearing mice (tumors were injected into the right liver lobe) were compared to tumor-free mice on day 15 (Figure 6X). On day 15, the left portion of the liver distal to the tumor site was harvested, and type 4 collagen levels were analyzed using RT-qPCR (Figure 6X). Liver tumor-bearing mice showed a 10-fold increase in type 4 collagen in the non-tumor liver (Figure 6Y). Additionally, when tumors were implanted in the subcutaneous space, an increase in collagen type 4 was found in the liver (Figure 6Y).

In summary, both local and distal tumor signaling correlated to the profibrotic polarization and enrichment of an IC population in the liver, as found through scRNA-seq of patient samples and qPCR of mouse livers.

## Discussion

We studied tumor-reactive CD8+ T cells using a high-dimensional, integrated approach utilizing three human scRNA-seq data sets (including appropriate per-patient analysis), human CODEX imaging, murine models, *in vitro* assays, and *ex vivo* systems in livers and primary tumors.^55^ Both in mice and patients, we found an unexpectedly high frequency of tumor-reactive CD8+ T cells in liver metastasis. Consistent with previous studies, we found that tumor-reactive CD8+ T cells displayed a dysfunctional state within the liver TME in both murine models and human scRNA-seq data.^28,29^ Our comprehensive analysis of the human scRNA-seq data indicated a variable phenotype of tumor-reactive CD8+ T cells across different organs. Furthermore, SPP1+ macrophages were found to interact with tumor-reactive CD8+ T cells in humans. These SPP1+ macrophages are distinct from the M2 macrophages *in vivo* and *in vitro* consistent with previous studies.^18^ Additionally, tumor signaling was seen to affect profibrotic polarization in the liver enriching the SPP1 interactions and contributing to CD8+ T cell dysfunction in the tumor microenvironment. This signaling enriched an IC population that was found to connect KC and SPP1 macrophage populations using pseudotime trajectory analysis and positively regulate collagen production. Furthermore, analysis of the healthy liver of subcutaneous tumor-bearing mice revealed elevated collagen production.

The presence of SPP1+ macrophages correlates with worse survival in various cancer types.^18,38,39^ However, our understanding of their biological function in the TME and their effects on tumor-specific T cell responses remains limited.^18,20,24^ For decades, M2 macrophages have been recognized as cells that promote tumor growth.^56,57^ Both functional murine *in vivo* studies and comprehensive bioinformatic analysis of human scRNA-seq data on T cells and M2 macrophages have provided findings complementary to the protumor role of SPP1+ macrophages.^10,17^ Previous studies have shown the potential role of SPP1+ macrophages on effector T cells, hypothetically separate from the M1/M2 dogma^18^; however, there is limited understanding of whether SPP1+ macrophages can induce tumor-reactive CD8+ T cell exhaustion and dysfunction without an enriched M2 macrophage population.^18–21,23^ Previous studies on both T cell graveyard and SPP1 interactions have focused on intrahepatic mediators. The T cell graveyard or responder trap phenomenon describes an accumulation of apoptotic, activated CD8+ T cells in the liver.^6,7^ This mechanism is mediated by Kupfer cell (KC) expression of FasL & nitric oxide, CD8+ T cell upregulation of LFA-1, antigen presentation by liver sinusoidal endothelial cells (LSEC) & hepatocyte and intrahepatic expression of IL-2, IL-4, IL-12, IL-15 and IL-18.^6,7^ In contrast, we describe a tumor- and liver-specific mechanism where extrahepatic tumor signals cause the overexpression of an intermediate macrophage cell population. This intermediate macrophage population then upregulates collagen deposition and enriches immunosuppressive SPP1+ macrophage-CD8+ T cell interactions.

SPP1 macrophages have been found to interact with fibroblasts in the colon and CRC metastasis in the liver.^19,20^ SPP1 ligand and PD-L1 on SPP1 macrophages promote CD8+ T cell exhaustion in vitro as well as in lung adenocarcinoma and gastric cancer.^21–23^ Recent studies have found SPP1 macrophages to be a distinct macrophage subset from KCs in fatty liver.^24^ Rather than a SPP1 population expanding only after seeding of metastatic cancer cells, we postulate a mechanism linking primary extrahepatic tumors to premetastatic expansion of a intrahepatic TGF-βR+ IL6R+ SPP1-KC intermediate macrophage population.

Previous literature found in fatty liver that these SPP1 macrophages were distinct from KC and were derived from monocytes.^24^ This study provides insight into their role within the liver TME including a unique mechanism of their development. Here, we find that SPP1 macrophages were KC-derived through TGF-B and IL6 signaling. Additionally, IC enrichment in the premetastatic niche may predict liver metastasis by promoting tumor-specific CD8+ T cell exhaustion and enrichment of fibrosis in the TME.

Multi-omic scRNA-seq-CODEX analysis of livers indicated that SPP1+ macrophages and CD8+ T cells interact in a profibrotic liver environment. Time course experiments revealed CD8+ T cell exhaustion in the liver of mice with subcutaneous tumors prior to tumor metastasis. This finding led us to hypothesize that primary tumors may promote the formation of profibrotic areas in the liver and explain the preferential formation of metastasis in the liver. Previous studies have found key chemokines and ligands involved in both metastasis of other cancer types as well as liver fibrosis.^58–60^ Specifically, TGF-beta and IL-6 are one of the few cytokines overlapping in both fields of study.^58–60^ TGF-β has previously been found in both prostate cancer and breast cancer to induce distant bone metastasis by establishing driver gene mutations that create a premetastatic niche in distant tissue.^58,59^ Similarly, IL-6 was identified as a mediator for cross-talk between bone marrow and cancer cells in breast cancer models. Levels of IL-6 correlated with increased monocyte dendritic progenitor growth along with an increased incidence of metastasis.^61^ TLR-4 potentiates TGF-β production and action while TLR-2 functions similarly to IL-6.^48–51^ Our data suggests that these profibrotic proteins are signaling to the distant liver enriching an intermediate cluster correlating to fibronectin and SPP1 expression and inducing CD8+ T cell exhaustion and dysfunction.

The size of the of human samples used in our scRNA-seq analysis was relatively modest and stratification based on survival for these samples was not possible as such data is not available at this time. However, three distinct scRNA-seq data sets were used in this analysis to further address these issues as best possible. Additionally, further research is needed on the effect of immune checkpoint blockade on the SPP1+ macrophage-CD8+ T cell interactions. Finally, this study had limited mechanistic data on the relationship between the profibrotic proteins and liver fibrosis since these specific mechanisms are discussed elsewhere in the literature including the large number of downstream secreted chemokines after receptor engagement.^51,62^

Future work is needed to study other organ sites enriched for SPP1+ macrophages-tumor-reactive CD8 T cells interactions. The SPP1+ macrophage-CD8 T cell interaction is a bidirectional targetable interaction, addressing an urgent need for a new avenue for potential drug development. Separately, we found changes to the fibrotic polarization in the liver TME causing tumor-reactive CD8 T cell dysfunction in advance of tumor spread. This result provides a secondary thread of interest for further research on the obvious clinical implications of reducing metastatic burden to the liver through reversal of this profibrotic machinery. In summary, here, we describe a novel mechanism by which primary tumors control distant CD8 T cells function by supporting the interaction of CD8+ T cells with SPP1+ macrophages in pro-fibrotic areas of the liver.

## Acknowledgements

RT and FJR-M receives funding through the NIH Medical Research Scholars Program. Research support was provided by the NIH Medical Research Scholars Program, a public-private partnership supported jointly by the NIH and contributions to the Foundation for the NIH from the American Association for Dental Research and the Colgate-Palmolive Company. TFG laboratory funding provided by the Intramural Research Program of the NIH, NCI (ZIA BC011345, ZO1 BC010870). TFG was support by NCI FLEX award.

## Data Availability

All data are available in the main text or the supplementary materials.

## Notes

### Competing Interest Statement

The authors have declared no competing interest.

