## Supplemental Figures for "A Paradoxical Tumor Antigen Specific Response in the Liver"

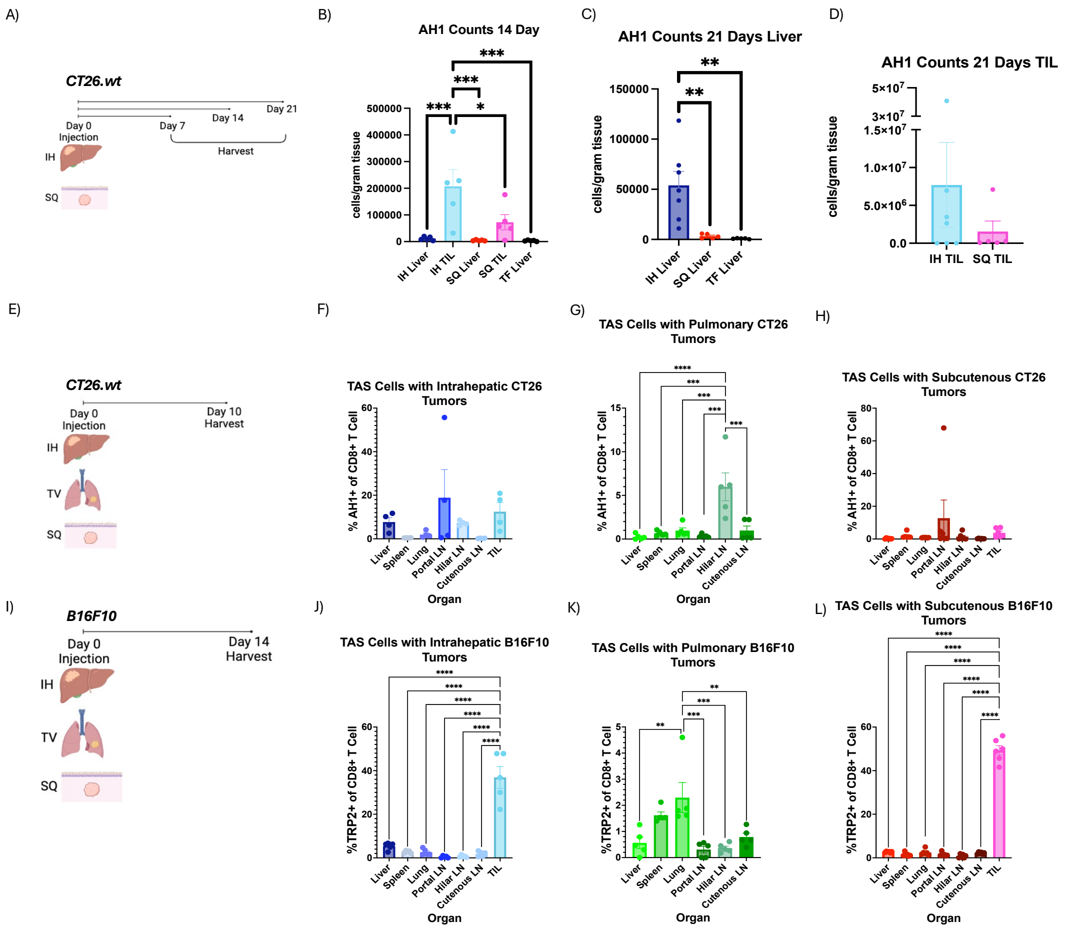


**Supplemental Figure 1. Organ Specific TAS CD8+ T cell Response. A)** Experimental schematic of B-D. Absolute counts of AH1+ CD8+ T cells at day 14 (n=5) (**B**), and at day 21 in the liver (IH, n=7; SQ, n=5) (**C**) and TILs (IH, n=7; SQ, n=5) (**D**) in CT26 tumor-bearing mice. **E)** Experimental schematic of F-H. Frequency of AH1+ CD8+ T cells in liver (n=4) (**F**), pulmonary (n=5) (**G**), and subcutaneous (n=H) (**H**) CT26 tumor-bearing mice. **I)** Experimental schematic of J-L. Frequency of AH1+ CD8+ T cells in liver (n=5) (**J**), pulmonary (n=5) (**K**), and subcutaneous (n=6) (**L**) B16F10 tumor-bearing mice. Data was analyzed by ordinary one-way ANOVA (B-C, F-H,J-L).

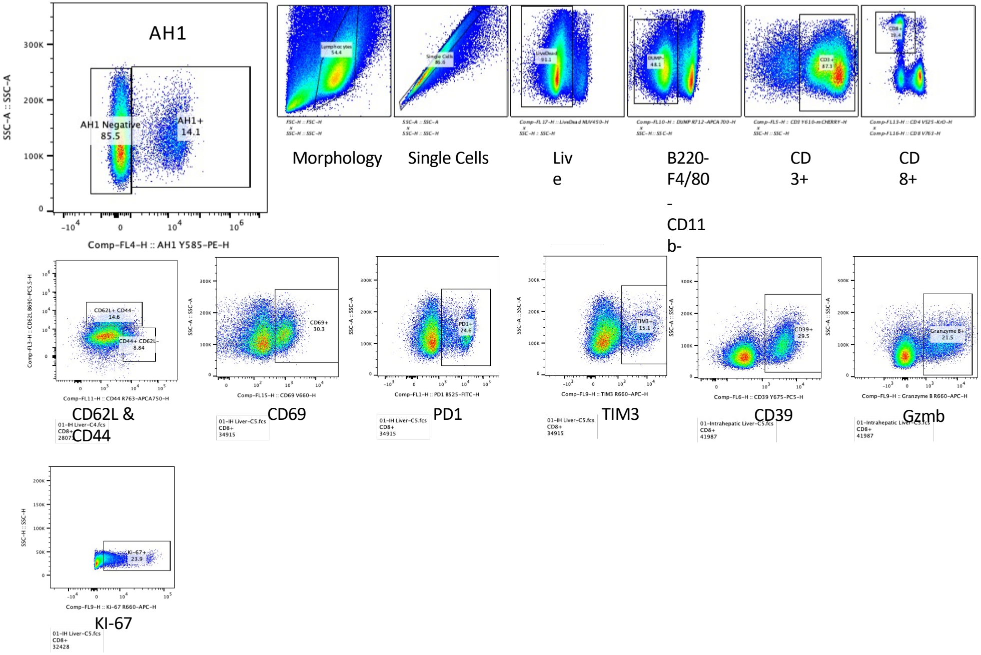


**Supplemental Figure 2. Flow gating strategy of TAS CD8+ T cells.**

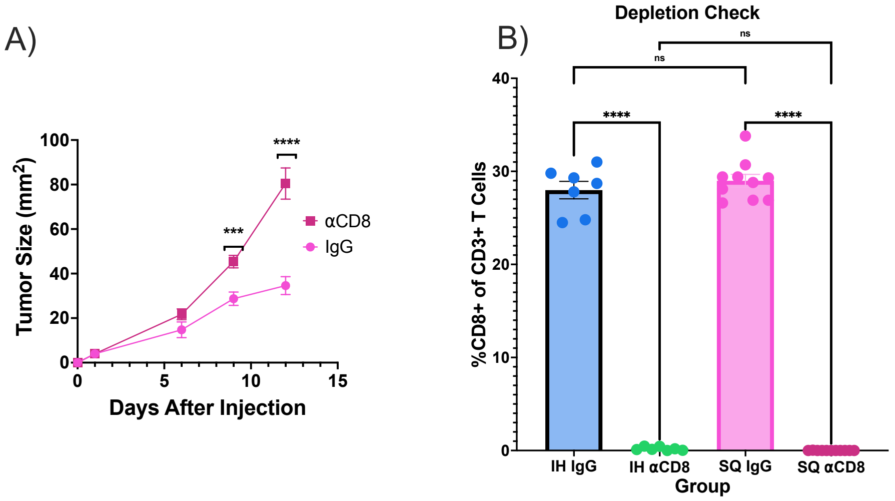


**Supplemental Figure 3. Effects of CD8+ T cell depletion on tumor growth. A)** Tumor size measurements after CD8+ T cell depletion (n=11) compared to IgG control (n=10) after subcutaneous injection. **B)** Flow cytometry staining of CD8+ T cells after CD8+ T cell depletion (IH, n=7;IgG,n=11) or IgG treatment (IH, n=7;IgG,n=10). Data was analyzed by unpaired t-test (A) and ordinary one-way ANOVA (B).

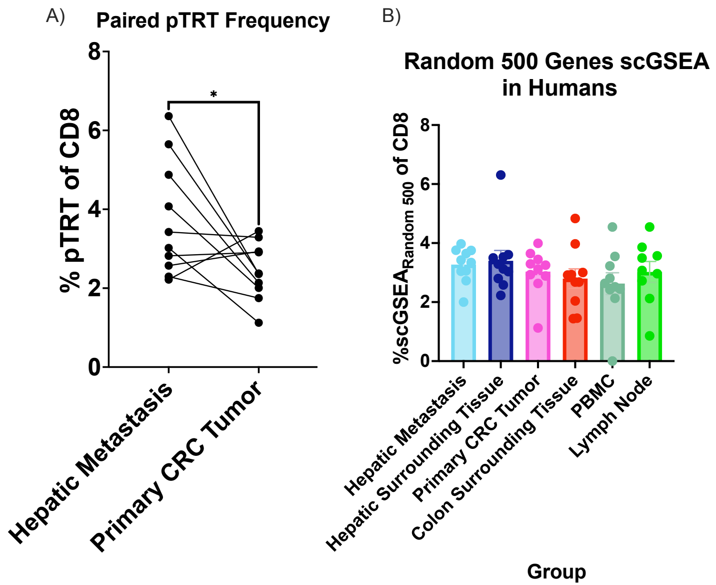


**Supplemental Figure 4. pTRT in human single cell analysis. A)** pTRT frequency of CD8+ T cells in hepatic metastasis compared to primary CRC shown as paired analysis (n=10). **B)** scGSEA analysis using same methodology as Figure 3A except with a random set of 500 genes (n=10). Data was analyzed by paired t test (A) and one way ANOVA (B).

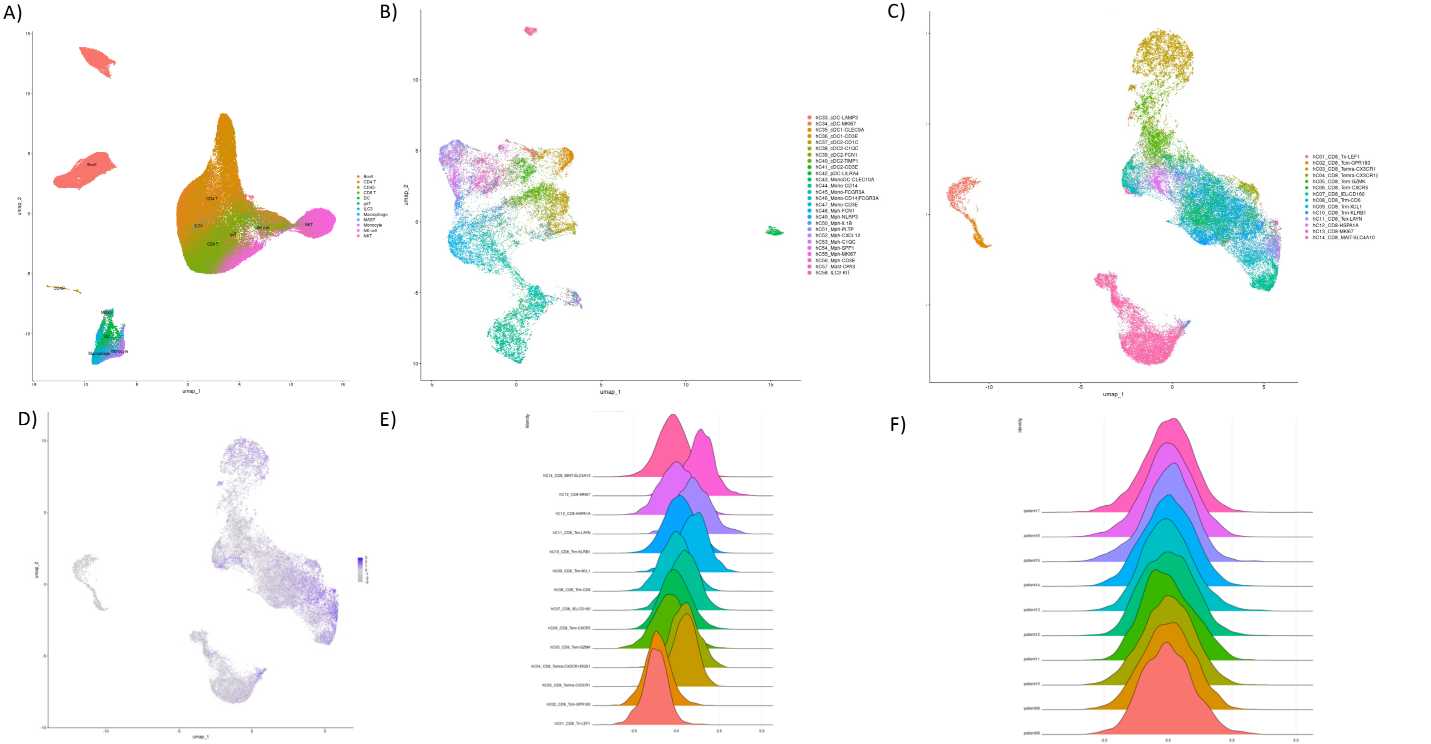


**Supplemental Figure 5. UMAP of CD8+ T cell clusters and pTRT scores.** UMAP showing **(A)** all cells, **(B)** myeloid clusters, **(C)** CD8+ T cell clusters and pTRT score based on scGSEA analysis as a **(D)** feature plot**, (E)** ridge plot per cluster, and **(F)** ridge plot per patient (n=10 patients).

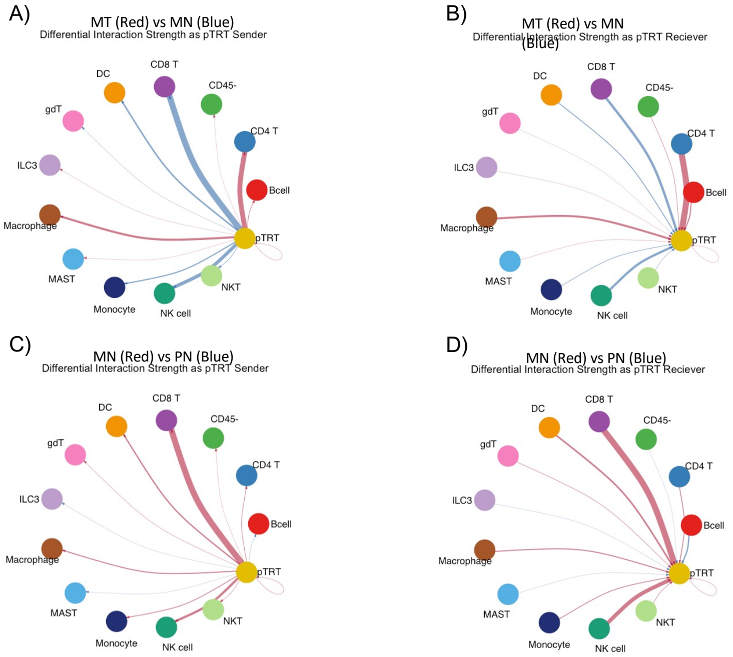


**Supplemental Figure 6. CellChat analysis of metastatic tumor, surrounding hepatic tissue and surrounding colonic tissue. A)** CellChat cellular communication analysis for communications that the pTRT are sending to all major cell types comparing the strength of interactions between hepatic metastasis (MT) to surrounding hepatic tissue (MN). **B)** CellChat cellular communication analysis for communications that the pTRT are receiving from all major cell types comparing hepatic metastasis to surrounding hepatic tissue. **C)** CellChat cellular communication analysis for communications that the pTRT are sending to all major cell types comparing the strength of interactions between surrounding hepatic tissue (MN) to surrounding colonic tissue (PN). **D)** CellChat cellular communication analysis for communications that the pTRT are receiving from all major cell types comparing the strength of interactions between surrounding hepatic tissue (MN) to surrounding colonic tissue (PN). Data was analyzed paired wilcox test (default for CellChat) (A-D).

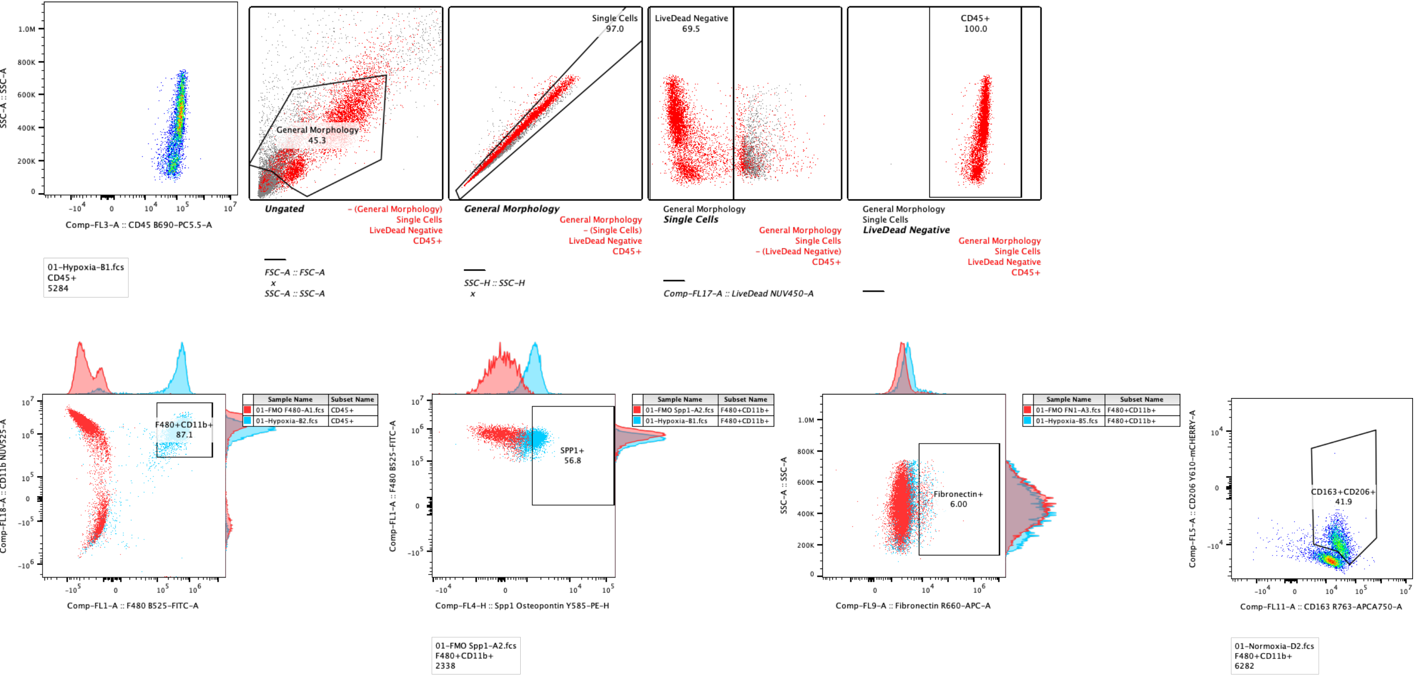


**Supplemental Figure 7. Flow gating strategy of macrophages.**

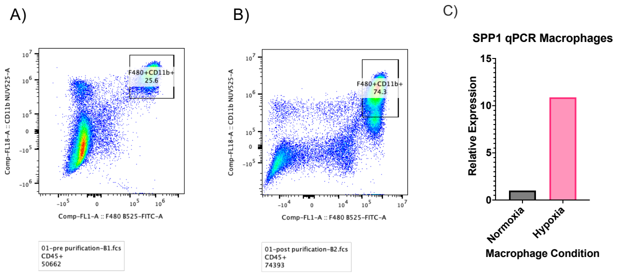


**Supplemental Figure 8**. **SPP1 macrophage isolation. A)** Peritoneal lymphocytes prior to sorting for F4/80+ macrophages. **B)** Peritoneal lymphocytes after sorting for F4/80+ macrophages. **C)** qPCR analysis of SPP1 in sorted macrophages in normoxic or hypoxic conditions.

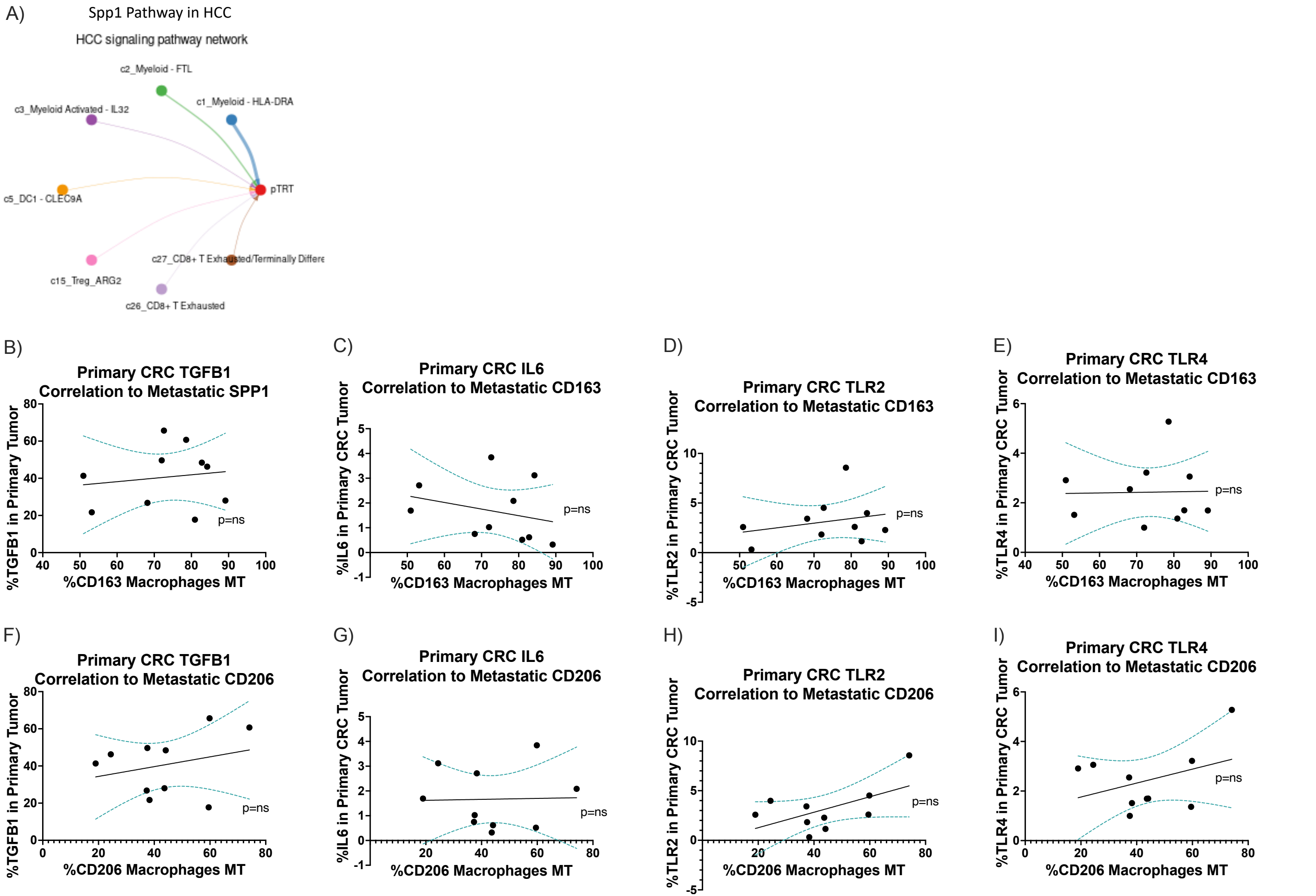


**Supplemental Figure 9. M2 Correlation in the setting of profibrotic polarization in the liver.** SPP1 CellChat pathway analysis for communication to pTRT in HCC scRNA-seq data (n=15) **(A)**. Correlation of metastatic intrahepatic tumor (MT) CD163 expression by macrophages to primary tumor TGF-β **(B)**, IL-6 **(C)**, TLR-2 **(D)**, TLR-4 **(E)** (n=10). Correlation of metastatic intrahepatic tumor (MT) CD206 expression by macrophages to primary tumor TGF-β **(F)**, IL-6 **(G)**, TLR-2 **(H)**, TLR-4 **(I)** (n=10). Data was analyzed using paired wilcox test (default for CellChat) (A) and simple linear regression (B-I).

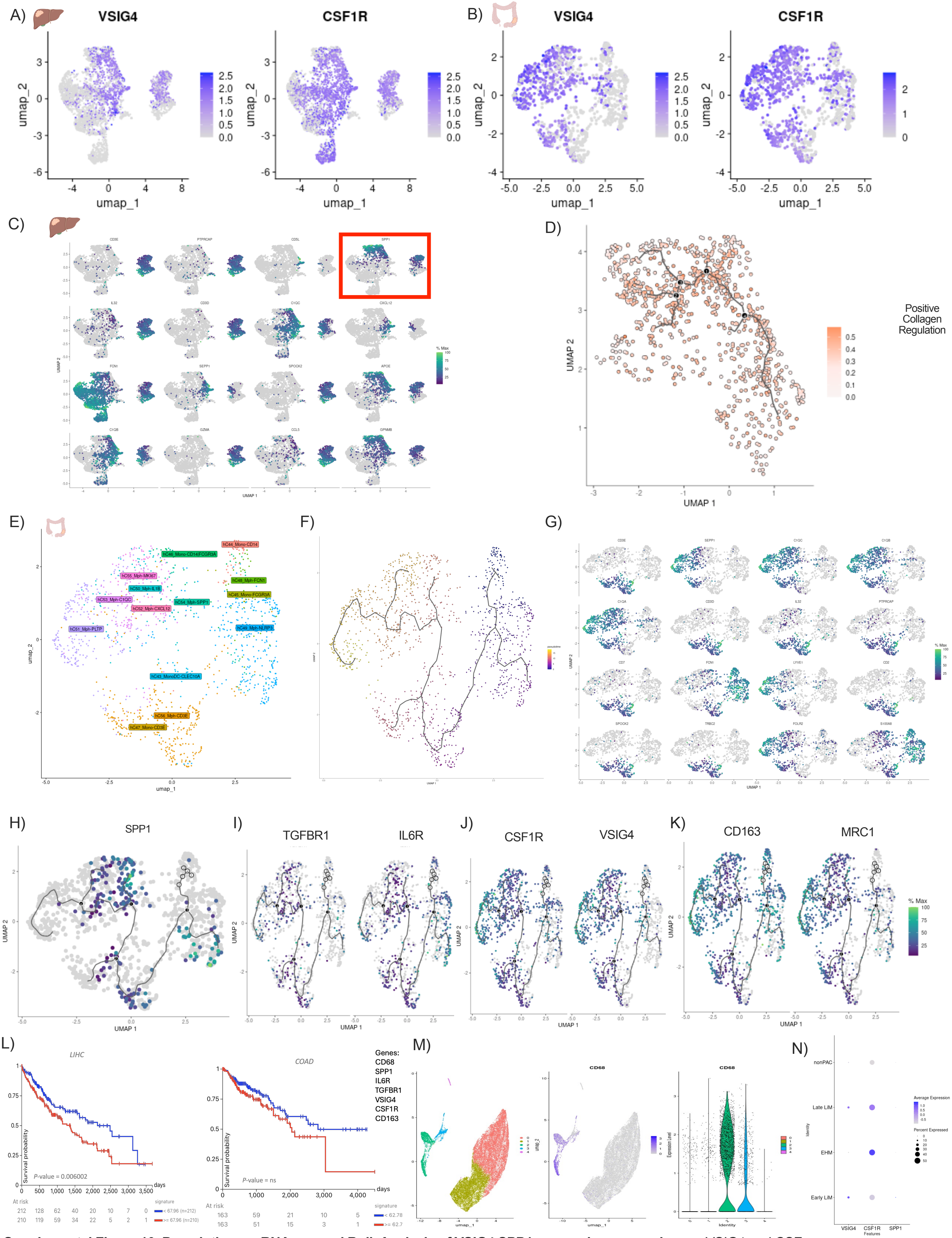


**Supplemental Figure 10. Pseudotime, scRNA seq and Bulk Analysis of VSIG4 SPP1 expressing macrophages.** VSIG4 and CSF1R feature plot of intrahepatic macrophage and monocyte population from metastatic CRC tumor-bearing liver **(A)** (n=10). VSIG4 and CSF1R feature plot of intracolonic macrophage and monocyte population (n=10) **(B).** Top genes in rank order correlating with liver pseudotime trajectory analysis (n=10) **(C)** with SPP1 boxed. Pseudotime trajectory analysis showing correlation of collagen regulation score (GO:0032967; positive regulation of collagen biosynthetic process) **(D)** with pseudotime trajectories in the liver (n=10). Intracolonic macrophage and monocyte population UMAP (n=10) from primary CRC tumor-bearing colon (**E**). Pseudotime trajectory analysis (n=10) of intracolonic macrophages and monocytes (**F**). Top genes correlating with colonic pseudotime trajectory analysis (n=10) **(C).** Pseudotime trajectory analysis showing correlation of SPP1 (**H**), TGFBR1 and IL6R (**I**), CSF1R and VSIG4 (**J**), and CD163 and CD206 (**K**) with pseudotime trajectories in the colon (n=10). Bulk TCGA survival analysis of HCC (left; n=438) and CRC (right; n=551) based on gene signature expression **(L).** UMAP (left) as well as feature plot (middle) and violin plot (right) of CD68 expression per cluster where cluster 2 and 3 were defined as CD68+ **(M)**. Kupfer cell and SPP1 marker expression per clinical group **(N)**. Data was analyzed using Morans I (C and D, G-K) and p-value (chi-squared distribution) (L).
